# On the Edge: Identifying priority areas for conservation of Fishing Cat, a threatened wetland felid, amidst rapidly altering freshwater landscapes

**DOI:** 10.1101/2022.01.16.476498

**Authors:** Tiasa Adhya, Priyamvada Bagaria, Partha Dey, Vanessa Herranz Muñoz, Anya Avanthi Weerawardana Ratnayaka, Ashan Thudugala, N.A. Aravind, James G. Sanderson

**Author notes:** Corresponding author: Tiasa Adhya.

## Abstract

1. Freshwater ecosystems have been most severely impacted in the Anthropocene with 27% of its species threatened with extinction. Fishing Cat is a globally threatened South and South-east Asian wetland felid that is also a highly rated Evolutionarily Distinct and Globally Endangered (EDGE) species, i.e., it is a global priority for conservation and research. Being an understudied species, knowledge gaps exist on its basic ecology, such as distribution and niche.
2. To address this, ensemble species distribution modeling (ESDM) was used to clarify doubts on its potential distribution and niche. To provide a relatable current context, loss of suitable habitat to urbanization (2010-2020) was estimated by analyzing range-wide survey data with environmental and anthropogenic variables (night-time lights and land surface temperature as proxies for urbanization).
3. Wetlands (18.36%) and elevation (17.15%) are the most important variables determining the ecological niche of Fishing Cat. It was predicted to be mainly restricted to low-elevation (<111 m) wetlands in river basins of South and South-east Asia. An estimated 23.74% suitable habitat was lost to urbanization.
4. Incrementally building on the ESDM outputs, high priority movement corridors and landscape conservation units were identified.
5. South Asia holds the core of the global Fishing Cat population with two very important regions - Ganges Brahmaputra Basin and Indus Basin - sharing transboundary areas with highly suitable habitat and many priority conservation units. The former is strategic to maintaining connectivity between South and South-east Asian Fishing Cat populations while isolation effects in the latter need investigation. Coastal wetlands of South-east Asia, though severely impacted, are crucial for the felid’s persistence.
6. More than 90% of Fishing Cat’s potential range lies outside the protected area network. Here, the felid can be adopted as a flagship species to conserve rapidly degrading low-elevation wetlands within a socio-ecological framework by involving multiple stakeholders.

## **1.** Introduction

The alarming rates of species loss during the ongoing sixth mass extinction surpass pre-human background estimates and continue because of increasing anthropogenic threats (Barnosky et al., 2011; Tittensor et al., 2014; Burivalova, Butler & Wilcove, 2018). Freshwater ecosystems, which are lifelines to both society and diverse species assemblages, have been the most severely impacted by these unprecedented threats but remain under-prioritized in the conservation domain (Albert et al., 2020; Tickner et al., 2020). Occupying less than 1% of the Earth’s surface, these ecosystems provide habitat to one-third of the world’s vertebrate species and 10% of all species (Dudgeon 2019), yet freshwater biodiversity populations have experienced dramatic declines with wetlands being lost three times faster than forests (Gardner and Finlayson, 2018). Approximately 27% of freshwater species and 42% of the 70 wetland dependent mammals are threatened with extinction (Balian et al., 2008; Tickner et al., 2020). Increasing socio-economic development in productive lowland regions is a significant threat to freshwater ecosystems (Darwall and Freyhof, 2016). In clarion calls published recently, scientists have stressed on integrating modern technology with field based knowledge for tracking endangered species of high conservation value to develop global freshwater conservation strategies (Arthington, 2021).

Fishing Cat (*Prionailurus viverrinus*) is a member of the threatened freshwater mammalian guild (Veron et al., 2008). Moreover, it is a highly rated ‘Evolutionarily Distinct and Globally Endangered’ (EDGE) species (EDGE score > 4), i.e, a global priority for research and conservation, but remains understudied (Tensen, 2018). It is one among the only two felids to have morphological adaptations for a semi-aquatic hunting niche (MacDonald and Loveridge, 2010), despite modern felids being morphologically and ecologically similar due to their recent and rapid divergence (Johnson et al. 2006). These include a double-coated, water-resistant fur, partially webbed feet, a short and stubby tail that provides balance while hunting or navigating in water and half-sheathed claws that provide traction in mud and facilitates gripping of slippery prey like fish (Kitchener, 1991; Sunquist and Sunquist, 2014; Hunter, 2019). Further, fish is known to be an important prey species and the primary constituent of its eclectic dietary niche (Haque and Vijayan, 1993; Cutter, 2015; Mukherjee et al., 2016). Both its morphological adaptations and prey association therefore suggests a strong selection for thriving in wetland environments.

Confirmed records of its presence exist from Pakistan, India, Nepal, Bangladesh, Sri Lanka, Myanmar, Thailand, Cambodia, Vietnam and Java (Willcox, 2015; Mukherjee et al., 2016, Lin and Platt, 2019) while its presence in peninsular Malaysia, Sumatra, Lao PDR, Taiwan and China remains speculative at best (Duckworth et al., 2009; Sanderson, 2009; Jutzeler, 2010; Duckworth et al., 2010). Further, there is lack of clarity on its distribution even within range countries. For instance, was/is it present in India’s western coast? Though there is anecdotal evidence and speculation of its occurrence there (Jerdon, 1874; Nowell and Jackson, 1996; Sunquist and Sunquist, 2014), recent surveys have failed to detect it (Janardhanan et al., 2014). Its actual distribution in South-east Asia still needs clarity with targeted research being conducted only in Thailand and Cambodia (Cutter, 2015; Thaung et al., 2018; Chutipong et al., 2019). Moreover, recent records of its occurrence from relatively non-moist drier areas (Sadhu and Reddy, 2013; Talegaonkar et al., 2018) and higher altitudes (>1800 m) (Thudugala, 2015) further create confusion regarding limitations of the species’ fundamental niche.

The tools and techniques offered by Geographical Information Systems (GIS) have immense potential in combining large scale ground information with remotely sensed satellite data and modeled spatial data layers to demarcate species’ geographical ranges, identify its movement corridors (Jalkanen et al., 2020; Xiao et al., 2020), and pinch points of connectivity (Yu et al., 2021), leading to prioritization of regions for their conservation (Lin et al., 2020; Shrestha et al., 2021). Species distribution modeling (SDM) approach has demonstrated the applicability of machine learning in defining species-environment relationship and the geographic extent of species distribution (Ahmed et al., 2021; De Simone et al., 2021; Lozano, 2021). SDMs combine the theory of ecological niche (Hutchinson, 1957), with machine learning algorithms to estimate a set of conditions appropriate for the species, and thereby estimate its potential spatial extent of distribution. These predictive models combine species presence locations with environmental variables (Araujo and Guisan, 2006; De Simone et al., 2021) to not only estimate species’ geographic extent but also identify the limiting environmental factors. However, performance of these predictive models may vary from case to case (Hao et al., 2019). Instead of selecting an individual algorithm, an ensemble of models harnesses better predictive power (Araujo and New, 2007; Hao et al., 2019; Lozano, 2021).

In the present study, we employed ensemble species distribution modeling (ESDM) to a) estimate the potential global distribution of Fishing Cat and b) identify factors limiting its fundamental niche. To understand the current extent of its distribution under the impact of urbanization, the most prominent marker of socio-economic development, we further c) estimated loss of suitable habitat to urbanization from 2010 to 2020. Incrementally building on the ESDM outputs, we identified d) critical movement routes in urbanized landscapes and e) Landscape Conservation Units (LCUs) to suggest conservation priority regions for maintaining global Fishing Cat habitat populations and connectivity.

## 2. Methods

### 2.1 Species Occurrences

Fishing Cat occurrences collected through field surveys by teams in respective Fishing Cat presence countries were collated (Appendix Data Sources). Only occurrences recorded through camera traps, or photographic evidence were used, while evidences that were not collected through systematic surveys (road kills, captures, dead specimens, museum specimens) were eliminated. Spatial filtering of the occurrence points was performed using a buffer of 4.6 km, assuming 20 km^2^ (∼4.6 km square grid) as the maximum home range for Fishing Cat (Sunquist and Sunquist 2014). The occurrences dataset was divided into temporal categories of those collected between 2010-2014 (T_1_) and 2015-2020 (T_2_). Here onwards we refer to the two time frames as T_1_ and T_2_. Records indicating road kill and conflict or negative interaction with humans were also retained for manual evaluation of model output.

### 2.2 Habitat Variables

Distribution models for Fishing Cat were developed accounting for two scenarios, an ecological scenario and an urban impact scenario. A set of 37 variables for the ecological scenario and two additional variables for the urban impact scenario were used (Appendix Data Sources). Values of the habitat variables were attached to the species occurrence locations using QGIS 3.10. Collinearity among the variables was computed using the R package ‘caret’ (Kuhn, 2008). Variables left after removing collinear variables (Fig 1 and 2), were also tested for Variance Inflation Factor (VIFs, Marquaridt, 1970). The variables retained were – annual mean temperature (bio_1), mean temperature of warmest quarter (bio_10), precipitation of driest month (bio_14), precipitation of warmest quarter (bio_18), mean monthly Normalised Difference Vegetation Index (NDVI) for February (NDVI_Feb), mean monthly NDVI for September (NDVI_Sept), elevation, ecoregions and wetlands (CIFOR). For the urban impact scenario, night-time lights and land surface temperature (LST) (Ariken et al., 2020) were used in addition to the ecological variables. To compare between the ecological and urban impact scenarios, we retained all the variables from the former in the latter scenario. The bioclimatic variables, elevation, eco-regions and wetlands remained constant across the two time intervals. NDVI, LST and night-time lights were used for the years 2012-13 and 2019 in order to represent the respective time intervals T_1_ and T_2_. Since the maximum home range for the Fishing Cat (20 km^2^) was considered, bioclimatic layers of spatial resolution 2.5 minutes (∼4.6 km) were selected and the other raster layers were resampled to 2.5 minutes spatial resolution. Considering the geographic scale of this study, all analyses were performed using Asia Lambert Conformal Conic meter projection.

**Fig 1:**
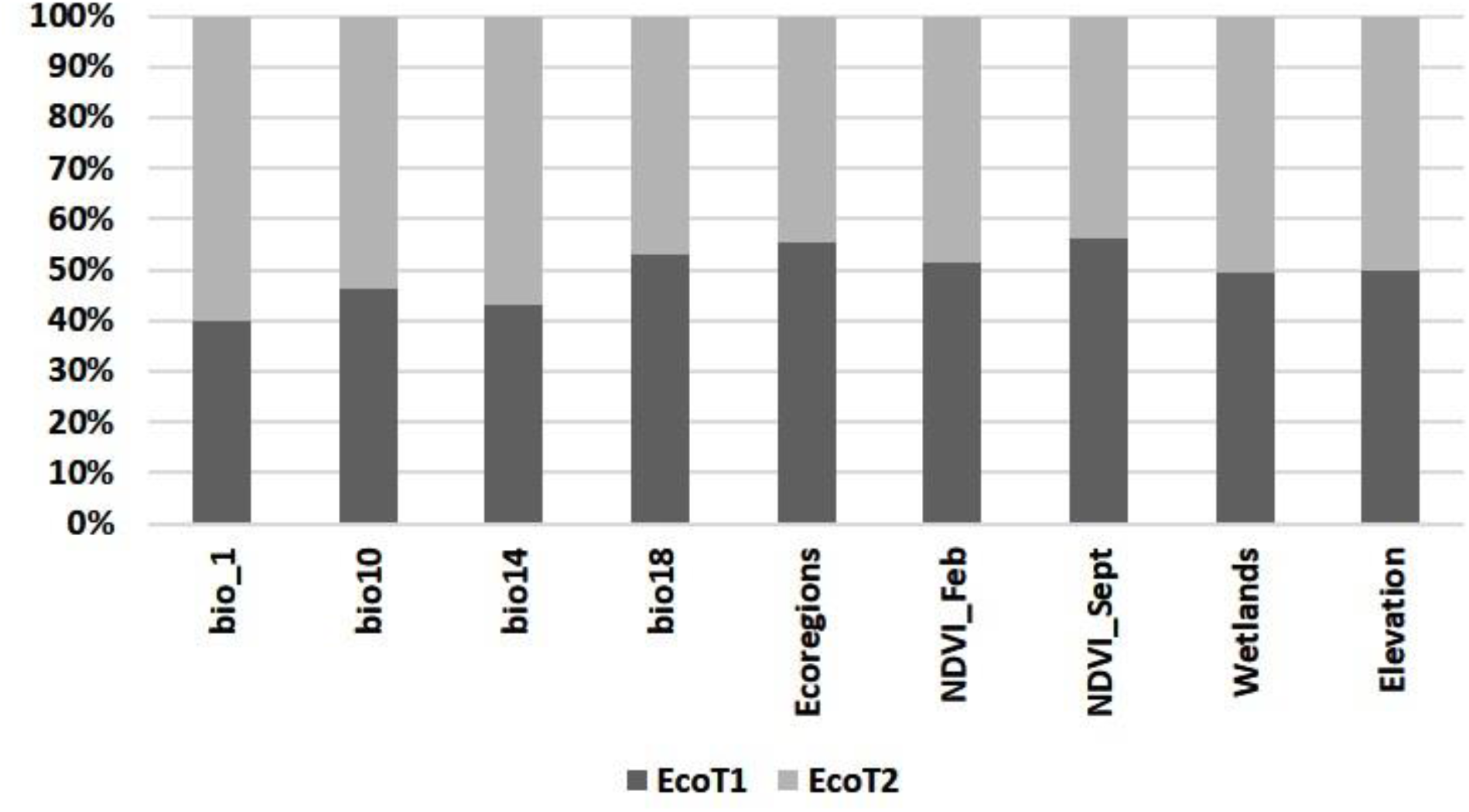
Variable Importance Scores for the ecological scenario.

**Fig 2:**
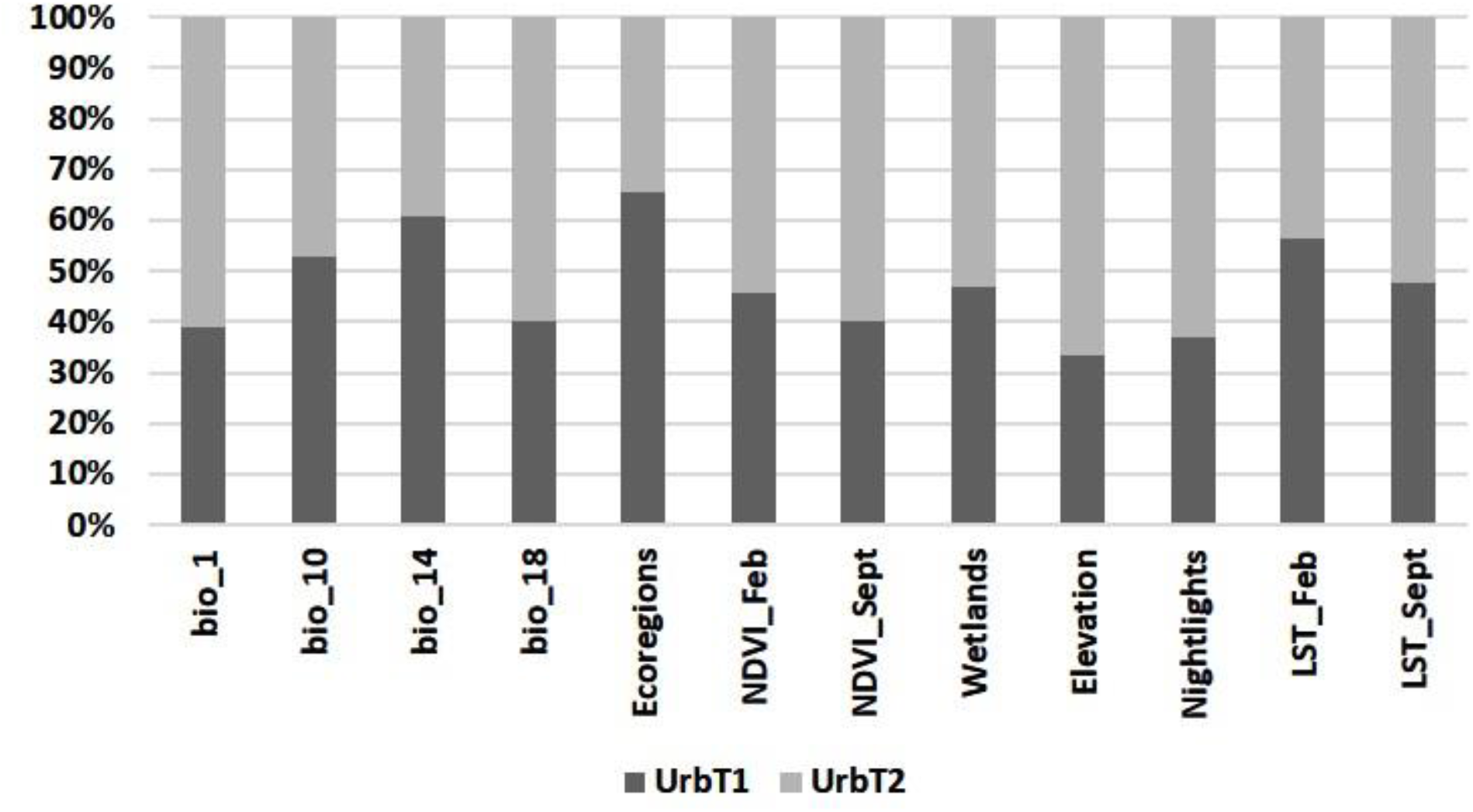
Variable Importance Scores for the urban scenario Bio 1 = Annual Mean Temperature. Bio 14 = Precipitation of Driest Month Bio 18 = Precipitation of Warmest Quarter Bio 10 = Mean Temperature of Warmest Quarter CIFOR = global wetland maps LST_Feb = Land Surface Temperature in February LST_Sept = Land Surface Temperature in September NDVI_Feb = Normalized Difference Vegetation Index for February NDVI_Sept = Normalized Difference Vegetation Index for September SRTM = global elevation data

### 2.3 Modeling Procedure

The ecological and urban impact scenarios were modeled for both T_1_ and T_2_. Hence, four different distribution models were developed representing the ecological scenario for T_1_ (EcoT_1_), ecological scenario for T_2_ (EcoT_2_); the urban scenario for T_1_ (UrbT_1_), and the urban scenario for T_2_ (UrbT_2_).

We used the ensemble modeling approach for developing the ensemble species distribution model (ESDM) for FC, using r package ‘SSDM’ (Schmitt et al., 2017). ‘SSDM’ is a robust and user-friendly platform for modeling ecological niche of the species, using mathematical representation of the known occurrences of the species (Guissan and Thuiller, 2005; Schmitt et al., 2017). The ensemble modeling approach has been proven to be useful in eliminating bias arising from single models and predicting more precisely (Araujo and New, 2007; Hao et al., 2019).

The model was supplied with a presence only occurrence records and calibrated to pick pseudo-absence/background points using the default strategy (Barbet-Massin et al., 2012) incorporated within the ‘SSDM’ package. Each algorithm was set for 15 replicate repetitions and ‘leave-out-one’ cross-validation strategy was used for evaluation of the ESDM. Receiver’s Operating Curve (ROC) was used for model evaluation and binary map threshold computation. Algorithm selection in ‘SSDM’ is automated. The algorithms which have an evaluation above a specified threshold, are only used in the final ensemble projection. A threshold ROC of 0.75 was set for algorithm selection. The algorithms which passed this filter were Random forest (RF), Support vector machine (SVM), Multivariate adaptive regression splines (MARS) and Classification tree analysis (CTA). Based on the model evaluation scores (Table 1), we retained RF and SVM for the final model predictions. The metric used for computing the threshold for binary maps was sensitivity-specificity equality (Liu et al., 2005; Jiménez-Valverde and Lobo, 2007). Scenario EcoT_2_ was first parameterised using this procedure. Thereafter, EcoT_1_, UrbT_1_ and UrbT_2_ scenarios were computed, keeping the model parameters constant, for comparability and uniformity among model outcomes.

**Table 1:**
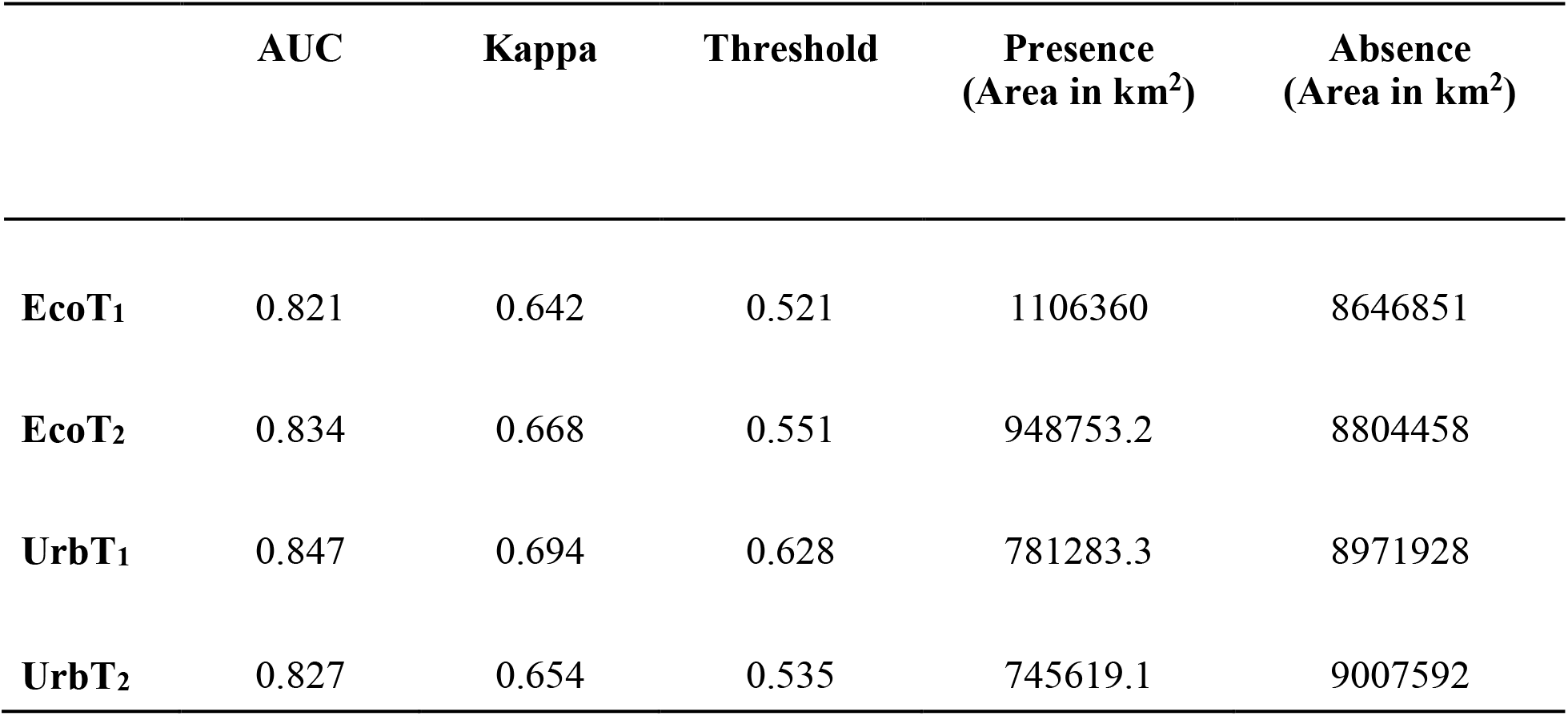
AUC values of all models along with threshold.

|  | AUC | Kappa | Threshold | Presence<br>(Area in km<sup>2</sup>) | Absence<br>(Area in km<sup>2</sup>) |
| --- | --- | --- | --- | --- | --- |
| **EcoT<sub>1</sub>** | 0.821 | 0.642 | 0.521 | 1106360 | 8646851 |
| **EcoT<sub>2</sub>** | 0.834 | 0.668 | 0.551 | 948753.2 | 8804458 |
| **UrbT<sub>1</sub>** | 0.847 | 0.694 | 0.628 | 781283.3 | 8971928 |
| **UrbT<sub>2</sub>** | 0.827 | 0.654 | 0.535 | 745619.1 | 9007592 |

Variable importance scores computation is automated in the ‘SSDM’ package. However, for variable response curves, random points were generated over the geographic range and values of all the variables used in the models and the probability surfaces were extracted to the points. The variable values were plotted against the probability scores using R package ‘ggplot2’ (Wickham, 2011). The limiting range of important variables determining Fishing Cat was estimated based on the occurrence data as was adopted by Mukherjee et al., 2010.

### 2.4 Loss of suitable habitat due to urbanization

The binary maps for the two scenarios and respective time intervals were used for calculation of suitable ecological niche for FC in the respective scenarios and the difference in the maps was calculated using the MOLUSCE 3.0.13 plugin in QGIS v2.18.2.

### 2.5 Habitat Connectivity

The ESDM based probability raster for UrbT_2_ was inverted to produce a friction layer for the movement of FC across its habitats. The ‘Least Cost Corridors and Paths’ module in the SDM toolbox (Brown, 2014; Brown et al., 2017) was used for identifying the corridors based on the least cost paths (LCPs) on a friction/cost layer. Due to processing limitations at the spatial resolution of 2.5 minutes (∼ 4.6 km), the friction layer for the LCP analysis using SDM toolbox was resampled to a resolution of 5 minutes (∼ 9.2 km). We used the ‘percentage of LCP value’ method for selecting the least cost corridors (LCCs) as this method selects corridors based on site-specific LCPs (Chan et al., 2011; Brown et al., 2017). Fishing Cat occurrences for T_2_ were used as input points. Due to the absence of any physical connection between Sri Lanka and the rest of the land area in the study area, and to avoid computational errors, Fishing Cat occurrence points from Sri Lanka were omitted from the LCC analysis.

In addition, a circuit theory-based connectivity analysis was also performed using the tool Circuitscape v4.0 (McRae et al., 2013), which implements the circuit and random walk theory (McRae et al., 2009). This tool computes a resistance-based connectivity metric based on a series of combinatorial and numerical analyses (Shah and McRae, 2008). The prime difference between a circuit and an LCP is that the circuit corridor is identified after computation of multiple paths between the population cores and the conductive probability of every raster cell is measured. In contrast, the LCP is based on a single least-cost route (Bowman et al., 2020). The circuit theory accommodates the assumption that animals may not have the knowledge of the perfect least-cost route (Marrotte et al., 2017) and the most traversed paths could be a cumulative outcome of the paths with minimum resistance. We used Protected Areas (PAs) where Fishing Cat occurrences for T_2_ were recorded as core areas indicating source populations. Pair-wise nodes and eight-cell window were parameterised for estimation of current flow, using the inversed ESDM as the resistance surface.

To pinpoint areas with very high density of both current flow and LCPs, i.e. the pinch points for Fishing Cat movement in the study region, the ‘Pinch Point Mapper’ tool (McRae, 2012) from the ‘Linkage Mapper’ toolkit for ArcMap (McRae and Kavanagh, 2011) was used. The Pinch Point mapper combines the frameworks of LCP and circuit theory and identifies pinch points having high density of paths represented through both the LCPs and Circuitscape based current flows. The pinch points are the bottlenecks in the corridor with considerable traffic of animal movement (McRae, 2012). These are regions with degraded or semi-degraded adjacent cells and thus points of significant importance for movement/dispersal from the landscape conservation perspective. Being narrow sections in the corridor, removal of these regions might hamper the connectivity of populations in a landscape (Liu et al., 2020). Linkage paths were constructed using the ‘Linkage Pathways’ tool, which were further used as inputs to the Pinch Point Mapper tool. Pairwise calculation was set for the Circuitscape mode and 4.6 km was set as the minimum cost-weighted corridor width. Since, all raster data in this study were processed at a spatial resolution of 2.5 minutes (∼ 4.6 km), this was the lowest possible corridor width that could be parameterised.

### 2.6 Priority Conservation Units

A conservation unit of 100 km^2^ was assumed for prioritization of landscape conservation units (LCUs) for Fishing Cat. The LCU prioritization was implemented through the R package ‘prioritizr’ (Hanson et al., 2018), using ‘Lsymphony’ open source solver. We used two different scenarios for modelling the LCUs – (a) costs based on FC connectivity and (b) costs based on urbanisation in the landscape. For the first scenario, the connectivity cost values from the LCP output raster and the inversed current flow raster were added and attached to each grid, to designate the costs that FC would incur in a given grid. These costs were calculated from the perspective of identifying the most suitable units for FC. For the urbanization scenario, the raster values from night-time lights and LST were combined and attached to the grids, representing the conservation costs that would be incurred if a given 100 km^2^ grid was to be selected for conservation. The raster output from the ESDM for UrbT_2_ and the CIFOR wetlands were used as feature layers.

Optimization targets ranging from 10%-50% of the landscape, with increments of 5% were implemented for each scenario resulting in nine solutions for each respective scenario. Prioritization importance scores (PIS) were assigned to each solution in the inverse order of their target optimization. A PIS of 50, 45, 40, 35, 30, 25, 20, 15 and 10 was assigned to solutions of targets 10%, 15%, 20%, 25%, 30%, 35%, 40%, 45% and 50% respectively (Rodewald et al., 2019). Hereafter, the solutions from all the nine prioritized targets were added into a single solution and units encompassing protected areas were selected.

## 3. Results

### 3.1 Global distribution of Fishing Cat and limiting factors

Wetlands (18.36%) and elevation (17.15%) were the highest contributing variables determining the ecological niche of Fishing Cat. The felid was predicted to be mainly restricted to low-elevation (<111 m above sea level) wetlands. These wetlands are part of the floodplains and delta regions of the Indus Basin, Ganges-Brahmaputra Basin (GBB), Ayeyarwaddy Basin, Salween Delta, Chao Phraya river system, Mekong Basin and Red River Delta (Fig. 3). In peninsular India, Fishing Cat was mainly predicted to occur in the Mahanadi Basin and the Godavari-Krishna Delta. The Lower Indus Basin and Middle Indus Basin constituting the westernmost limit of Fishing Cat’s distribution is predicted to have very high quality Fishing Cat habitat. Our model predicted the occurrence of Fishing Cat along the Indian western coast but the felid might have never occurred there (see Janardhan et al., 2014). Apart from this, the island country of Sri Lanka was predicted to have broad and continuous suitable areas for the cat along both the eastern and western coast. A narrow zone fringing the northern coastline of Java, another island, was also found to be suitable. Among these, the Indus Basin, parts of the Indo-Nepal Terai, Sunderbans delta along with low- elevational wetlands of the Gangetic floodplains, parts of the Mahanadi basin including Chilika lagoon, Bhitarkanika and Mahanadi mangroves and small portions of the Godavari- Krishna delta are predicted to have very highly suitable habitat. In South East Asia, such high quality habitats are sparingly present in the Ayeyarwady delta and coastlines of Thailand, Cambodia and Vietnam. More than 90% of the predicted suitable range for Fishing Cat lies outside protected areas (see Supplementary information, S1). To guide future surveys, we have provided detailed biogeographical information of regions with >50% predicted presence of Fishing Cat in the mainland and separately for the island country of Sri Lanka along with jurisdictional boundaries to maximize relevance for management (see Supplementary information, S2).

**Fig 3:**
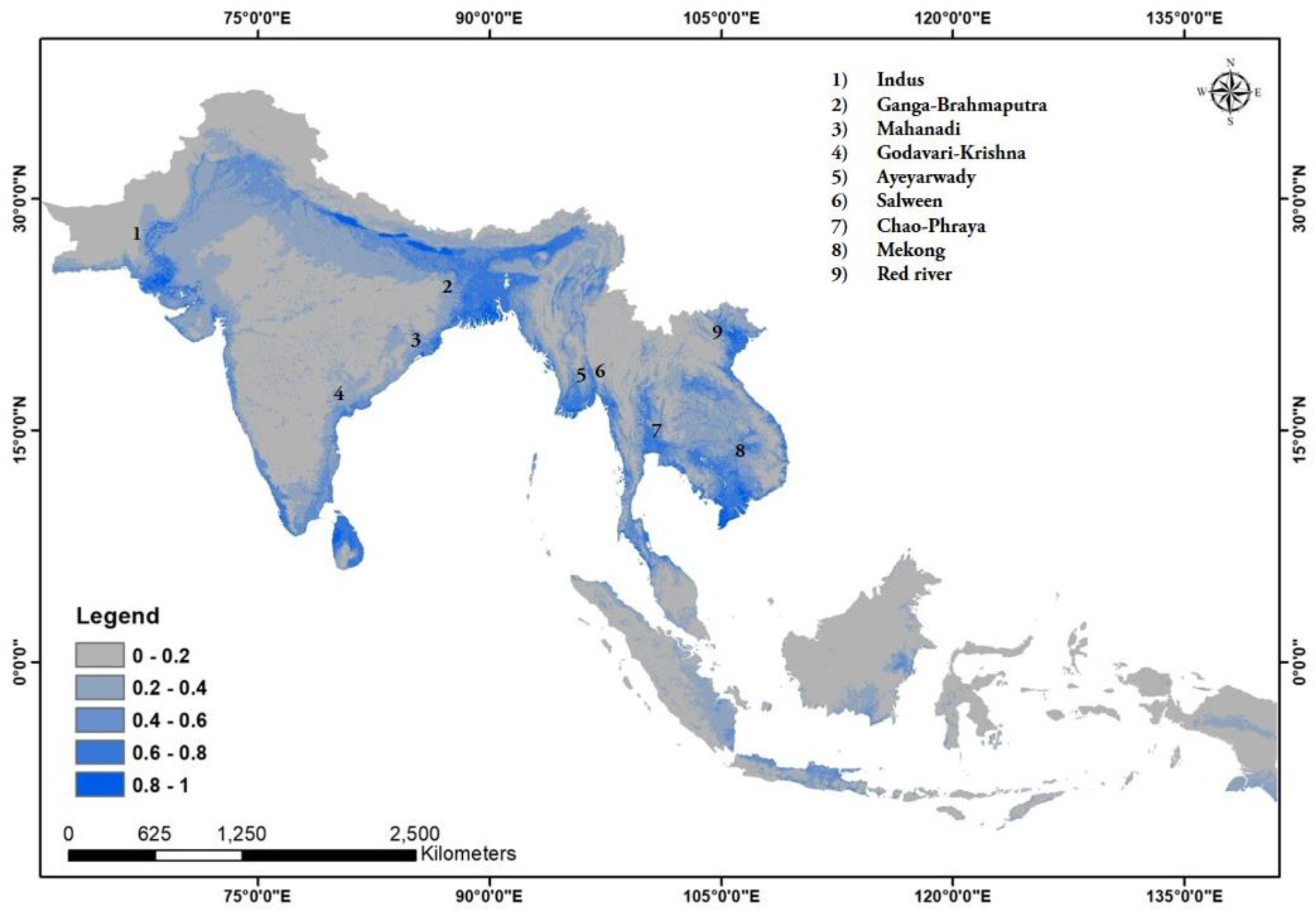
Predicted global distribution of Fishing Cat along river basins of South and Sout-east Asia with probabilities of occurrence.

**Fig 4:**
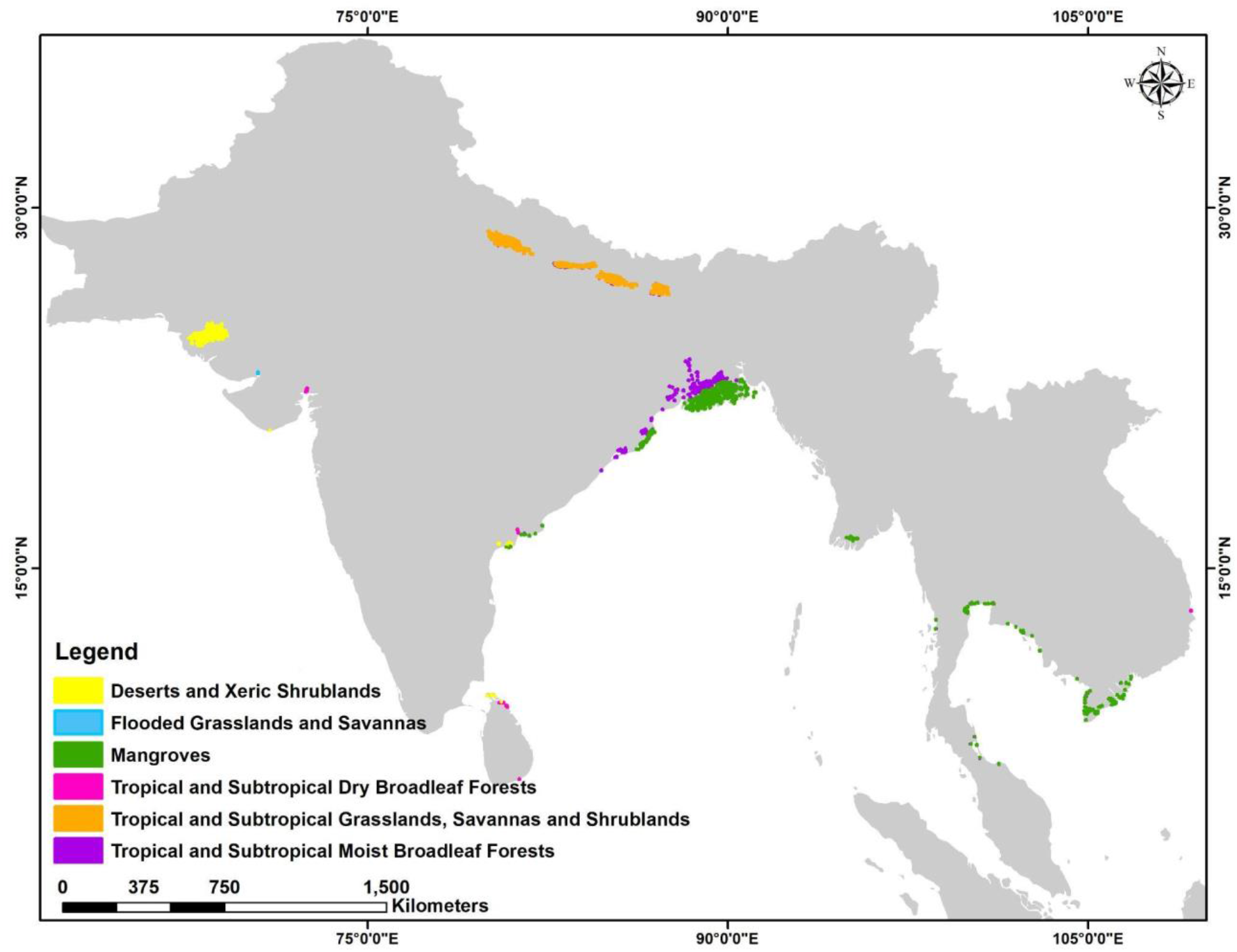
Ecoregions with Fishing Cat predicted to have more than 80% probability of occurrence. Note: Symbology inflated for better visualization.

### 3.2 Temporal loss of habitat

The ecological scenario (EcoT1) is a hypothetical one, useful in visualizing the extent of the suitable habitat for Fishing Cat with only natural predictors and without anthropogenic pressures. Since the bioclimatic and topographic factors remained constant over the two time points (T1 and T2) in this study, a decline of 18% (Table 2) in the suitable habitat under the ecological change scenario (EcoT1 to EcoT2) indicates a shrink in the vegetation cover/habitat, which indirectly represents anthropogenic pressure. On the other hand, the urban impact scenario (UrbT1 to UrbT2) includes proxies of anthropogenic structures as an influencing variable in the model and can better emphasize the impact of urbanization on Fishing Cat habitat.

**Table 2:** Changes in area of suitable habitat for FC in the two scenarios and time intervals. Note: Area in km^2^. EcoT_1_ - Ecological scenario for T_1;_ EcoT_2_ - Ecological scenario for T_2_; UrbT_1_ - Urban scenario for T_1;_ UrbT_2_ - Urban scenario for T_2_.

|  | EcoT<sub>1</sub> to EcoT<sub>2</sub> | UrbT<sub>1</sub> to UrbT<sub>2</sub> |
| --- | --- | --- |
| **Habitat gain** | 46014.0267 | 151373.806 |
| **Habitat loss** | 203621.1354 | 185824.7738 |
| **Percent gain (%)** | 4.159 | 19.375 |
| **Percent loss (%)** | 18.404 | 23.784 |

The ESDM probability maps that were generated using the selected algorithms, RF and SVM, had the Receivers Operating Curve (ROC) evaluation score above 0.82 and Kappa evaluation score above 0.64 for all the computed scenarios (Table 1). This indicates that we could successfully implement the models in estimating the suitable habitat for Fishing Cat in the two scenarios considered in this study. The binary ESDM maps were generated using the respective threshold values for each scenario (Table 1).

Area of the model predicted suitable area for Fishing Cat in each scenario was calculated (Table 1) and the changes in the predicted presence between T_1_ and T_2_ in both the ecological and urban impact scenarios were calculated (Table 2). The loss of predicted suitable area (Table 2) for Fishing Cat, in the ecological scenario (EcoT_1_ to EcoT_2_) was estimated at 18.404% and 23.784% in the urban scenario (UrbT_1_ to UrbT_2_) (see Fig 5 and 6). Simultaneously, percent gain in suitable habitat was also estimated, 4.159% for EcoT_1_ to EcoT_2_ and 19.375% for UrbT_1_ to UrbT_2_ (Table 2).

**Fig 5:**
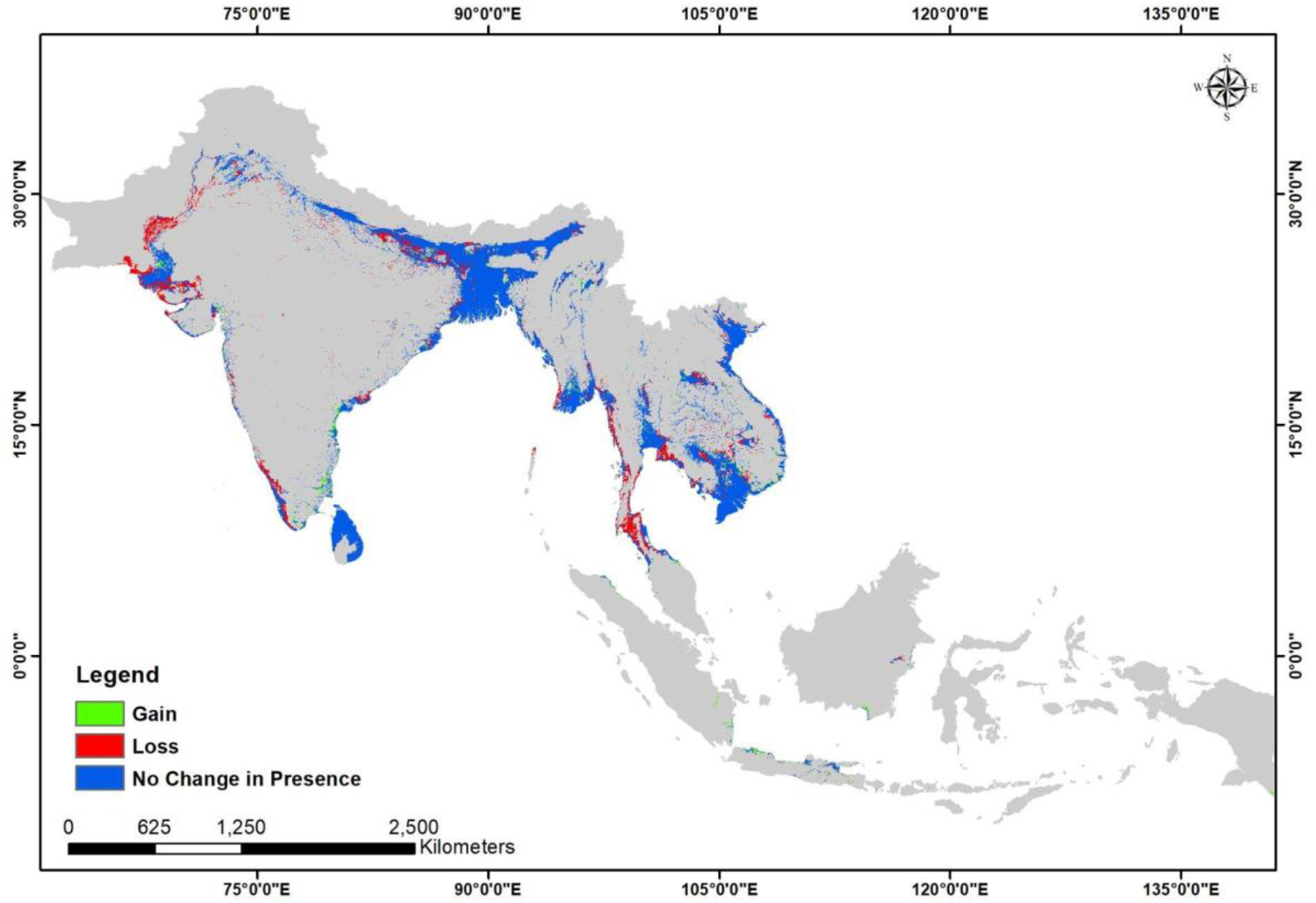
Change in the predicted distribution of Fishing Cat from 2010 to 2020, in an ecological change only scenario.

**Fig 6:**
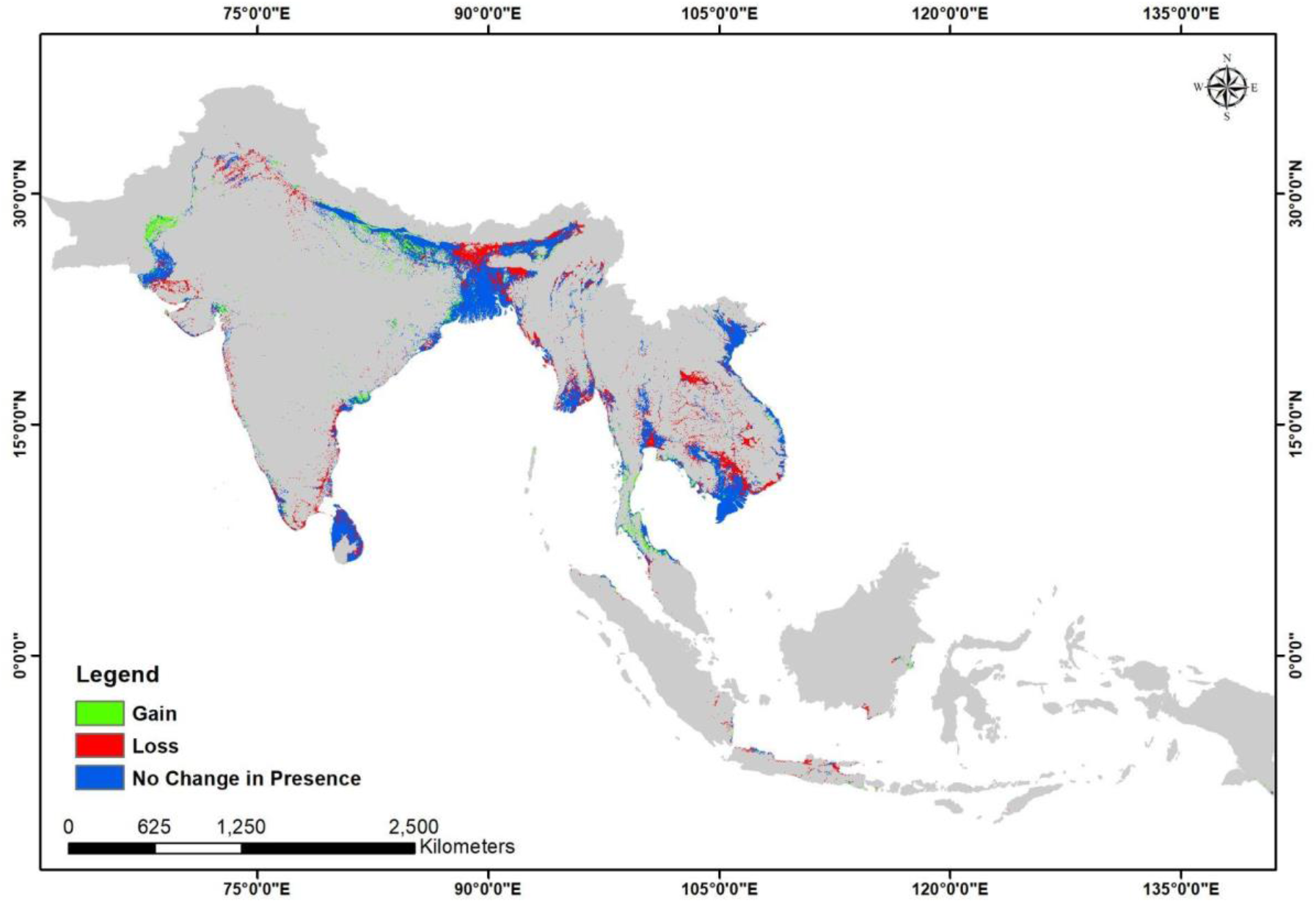
Change in predicted global distribution of Fishing Cat due to urbanisation between T1 and T2.

### 3.3 Landscape connectivity

The raster values of connectivity outputs i.e., the LCC, LCC line density; the Circuitscape output for cumulative current density and the Pinch-points were rescaled to 0-1. The lowest 30% of the cost values in the LCC were used to depict the LCC for Fishing Cat, and the highest 30% of the values to show the density of LCPs in the LCC (Fig. 7). Similarly, top 30% of the maximum flow values (Jennings et al., 2020; Li et al., 2020) were considered to depict the conductivity-based corridor for Fishing Cat movement (Fig. 8) and 5% of the highest values of the output from the Pinch Point Mapper were used as thresholds for delineating the pinch points (Fig. 9).

**Fig 7:**
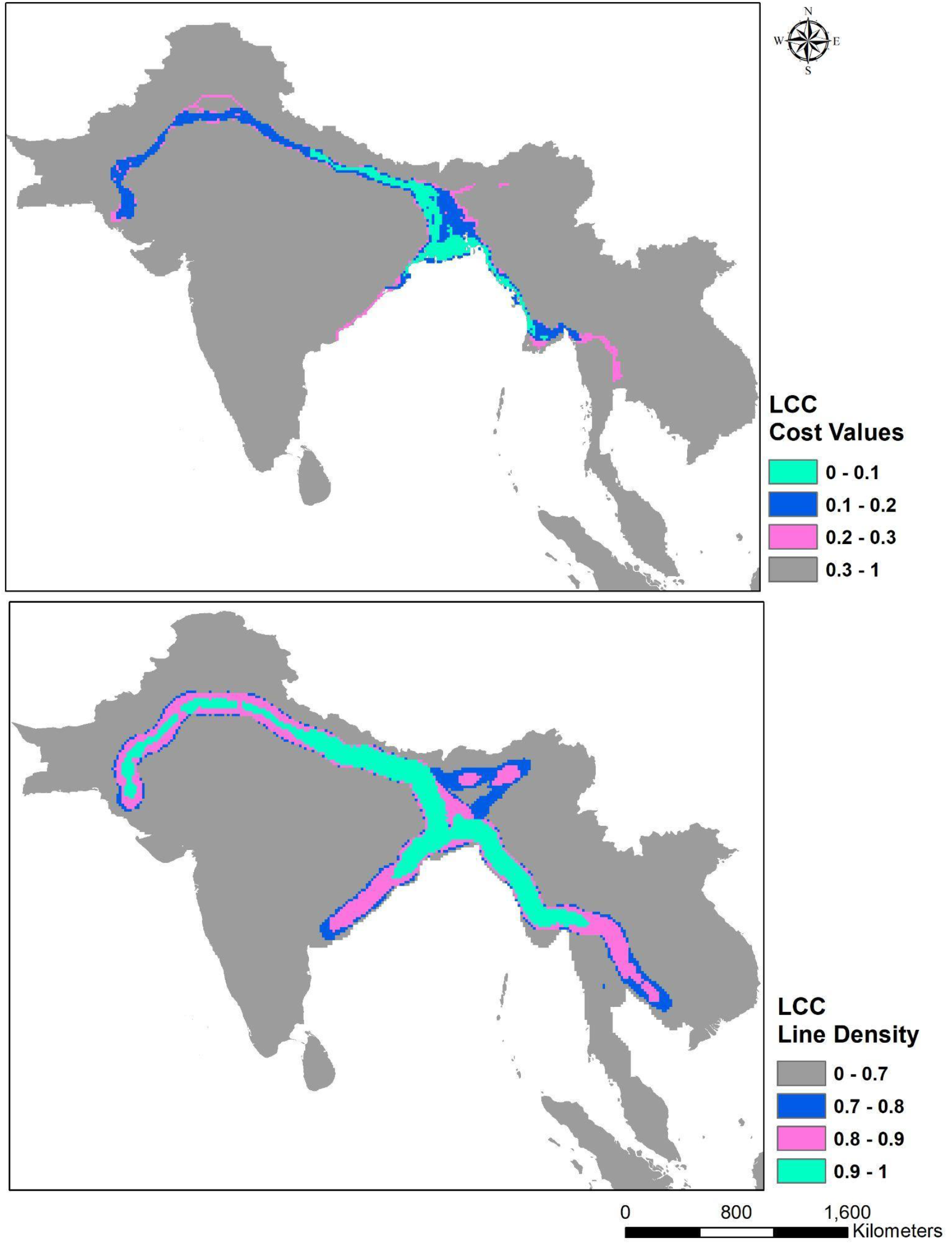
Least Cost Corridors (LCCs) and Line Densities of LCCs for global Fishing Cat connectivity. Note: Higher cost values in LCC denote higher friction in movement, while higher line density values denote most traveled LCCs.

**Fig 8:**
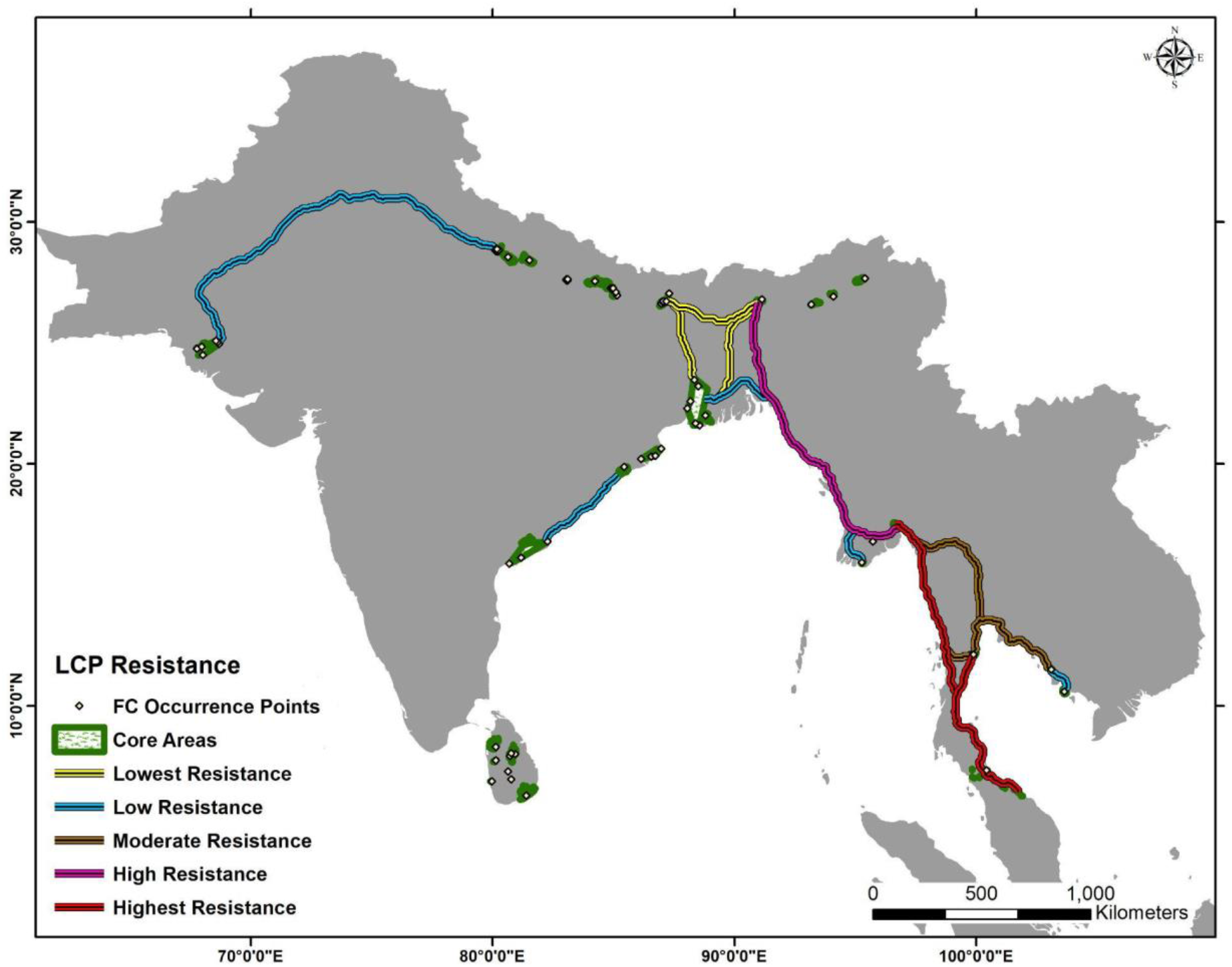
Least Cost Path (LCP) with varying levels of resistance to global Fishing Cat connectivity.

**Fig 9:**
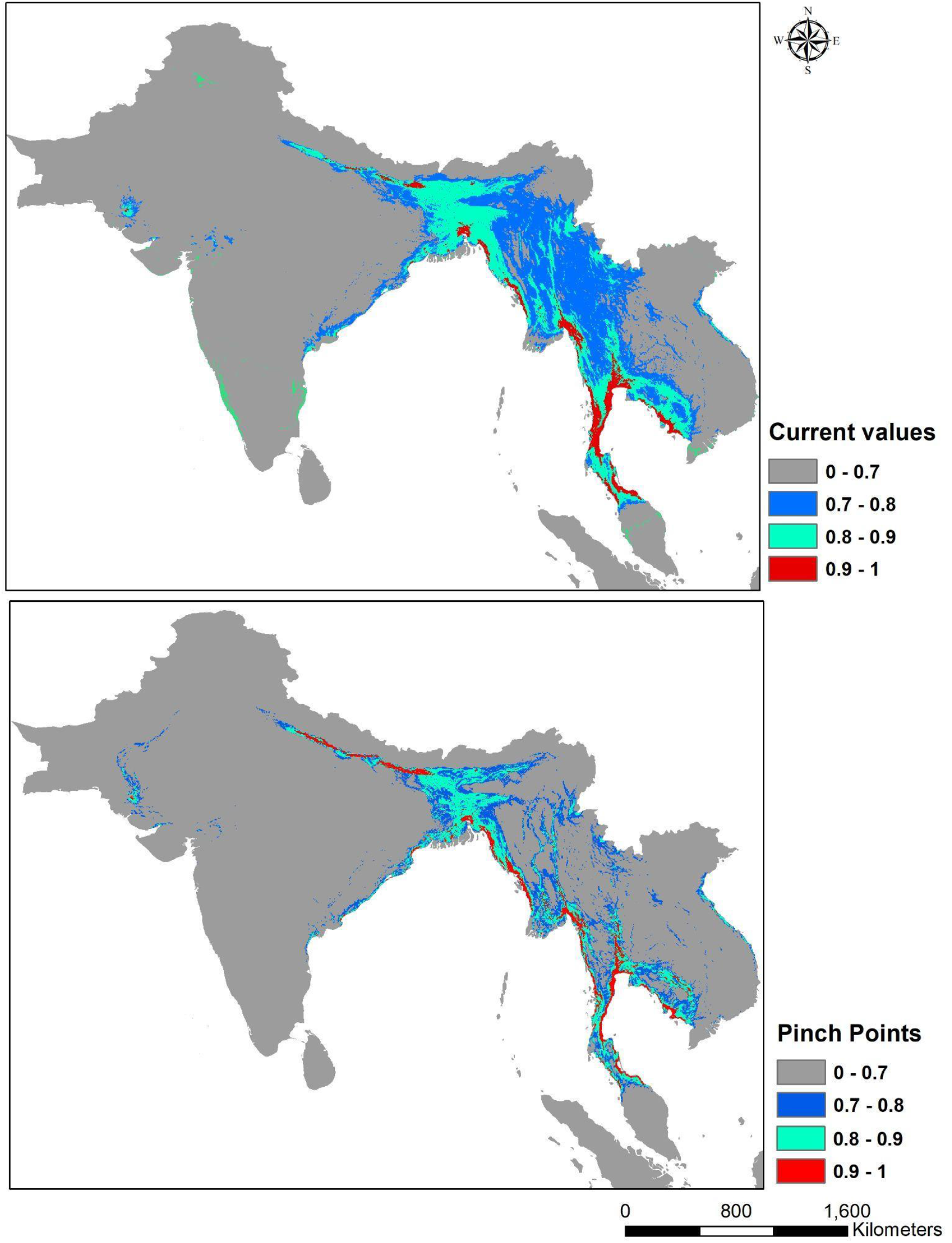
Circuitscape generated corridor showing higher current flow intensity in most suitable movement corridors (above). Corridors with a minimum width (4.6 km) and a high density of corridor path depict the pinch-points, i.e, bottlenecks in the corridor (below).

**Fig 10:**
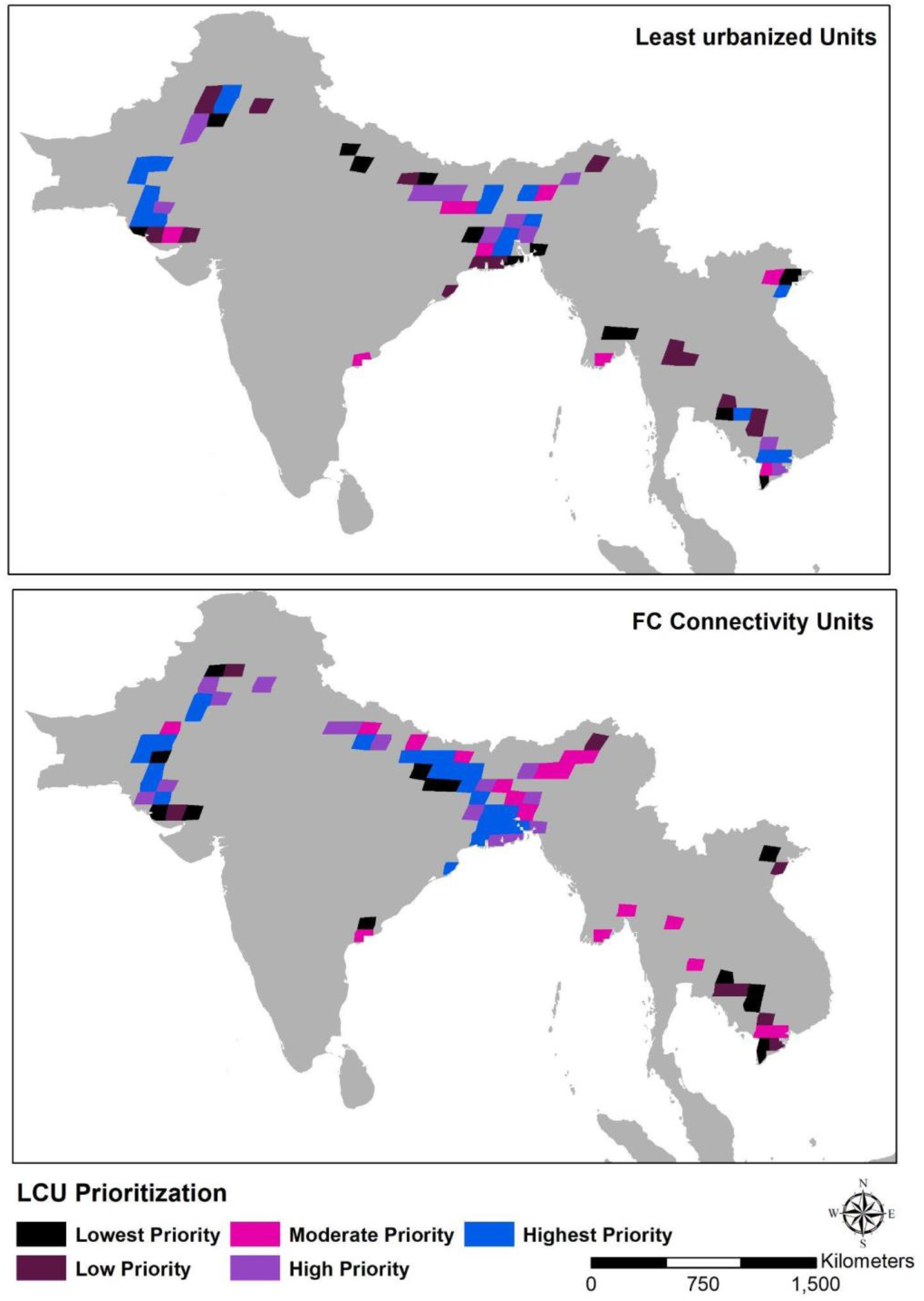
Landscape Conservation Units according to the least urbanized scenario (above) and most suitable habitat units that facilitate connectivity (below).

### 3.4 Landscape Conservation Units

In both the Fishing Cat connectivity scenario and the least urban scenario, the Indus basin and parts of GBB in South Asia figured as LCUs of very high priority.

According to the Fishing Cat connectivity scenario, 39 high to very high LCUs were found in South Asia. No high priority LCU was present in South-east Asia. Six LCUs of moderate priority were selected in South-east Asia. In the least urban scenario, 25 LCUs of high to very high priority were selected in South Asia. Six LCUs of high to very high priority were present in South-east Asia.

## 4. Discussion

To the best of our knowledge, this study was the first attempt to model the global distribution of Fishing Cat, an under-studied, globally endangered freshwater mammalian carnivore of high conservation and research value (Veron et al., 2008; Tensen, 2018; Zanin and Neves, 2019). RF and SVM, selected as the best performing algorithms for the SDM of Fishing Cat, have been found to be powerful classifiers in other studies too (Fukuda et al. 2013; Thahn Noi and Kappas, 2018). The felid was predicted to be mainly restricted to low-elevation (<111 m above sea level) wetlands of South and South East Asian river basins (Fig 3). Low elevation was also estimated to be an important predictor for Fishing Cat distribution in the Indian subcontinent (Silva et al., 2020). South Asia holds the core of the global Fishing Cat population with India sharing two trans- boundary regions in the GBB of high Fishing Cat occurrence probability (>80%) - a) wetlands of the Terai region with Nepal, and, b) the Gangetic Delta (including the Sunderbans) with Bangladesh. South Asia also has the highest numbers of LCUs in parts of the GBB and the Indus Basin (Fig 9, Table 3). The situation in South-east Asia seems precarious and therefore worthy of intensive research and conservation efforts. Along the coastline of Myanmar and Thailand, Fishing Cat connectivity is predicted to face the highest resistance but some areas of the Mekong floodplains and delta region provide lower resistance, higher habitat suitability including the presence of LCUs and potentially important dispersal corridors.

**Table 3:** Different LCUs selected in river basins of South and South-east Asia.

| **River Basins** | **Location** | **Highest Priority** | **High Priority** | **Moderate Priority** | **Total no of LCUs** |
| --- | --- | --- | --- | --- | --- |
| **Fishing Cat connectivity scenario** |  |  |  |  |  |
| Indus Basin | South Asia | 8 | 5 | 1 | **14** |
| GBB | South Asia | 15 | 10 | 10 | **35** |
| Mahanadi basin | South Asia | 1 | - | - | **1** |
| Godavari-Krishna delta | South Asia | - | - | 1 | **1** |
| Ayeyarwady basin | South-east Asia | - | - | 1 | **1** |
| Salween delta | South-east Asia | - | - | 1 | **1** |
| Chao Phraya river system | South-east Asia | - | - | 2 | **2** |
| Mekong Basin | South-east Asia | - | - | 2 | **2** |
| Red River Delta | South-east Asia | - | - | - | **0** |
| **Least urbanized unit scenario** |  |  |  |  |  |
| Indus Basin | South Asia | 9 | 3 | 1 | **13** |
| GBB | South Asia | 6 | 7 | 4 | **17** |
| Mahanadi basin | South Asia | - | - | - | **0** |
| Godavari-Krishna delta | South Asia | - | - | 1 | **1** |
| Ayeyarwady basin | South-east Asia | - | - | 1 | **1** |
| Salween delta | South-east Asia | - | - | - | **0** |
| Chao Phraya river system | South-east Asia | - | - | 2 | **2** |
| Mekong basin | South-east Asia | 3 | 2 | 1 | **6** |
| Red River Delta | South-east Asia | 1 | - | 1 | **2** |

### 4.1 Flexibility/limitedness of its fundamental niche

There are some departures from the hypothetical situation presented by our models as is expected, discussing which might resolve doubts and strengthen our understanding of its ecology and fundamental niche. For example, despite our predictions of the restriction of its niche to low-elevation wetlands (>111 m above sea level), Fishing Cats were documented from an elevation of 1870 m from Sri Lanka’s central hilly region. This is the only unit globally to colonize higher elevations and thus constituted a small fraction of the occurrence points in comparison to the regional species pool of the mainland. Therefore this was not chosen as a suitable habitat by our model. A closer introspection of the Sri Lankan geography reveals that freshwater sources in the island originate from the central hilly regions and flow out towards all directions, creating numerous water-rich habitat/wetland complexes throughout the island. Thus, during sea-level rise events, low-elevational populations might have been pushed to migrate up the hills due to geometric constraints imposed by the island and would have been facilitated to colonize the higher altitudes because of the presence of rivers. This could have caused a local adaptation in the Sri Lankan population and effectively extended its fundamental niche. Theoretical predictions do predict selection of broader niche breadth in species in variable environments (Levins, 1962; Sultan and Spencer, 2002). Mainland populations situated in more predictable environments may have been relaxed of such selection pressures. Additionally the presence of more competitors in the mainland might have limited them to a narrower and specialized niche. In addition, Fishing Cat was recorded from drier regions in Sri Lanka, also predicted to be unsuitable/sub-optimal by our model. The presence of man-made wetlands such as ancient hydrological tanks, reservoirs and croplands simulating Fishing Cat habitat might have facilitated its occurrence in these drier regions inland. Infact, annual rainfall was found to be the most important climatic variable determining Fishing Cat distribution in a Sri Lanka specific modeling (Thudugala, *in litt.*) that might be playing a critical role in maintaining these wetlands in the drier inland regions.

Fishing Cat has also been recorded recently from the much drier central and north-western Indian landscapes (Sadhu and Reddy, 2013; Talegaonkar et al., 2018; Dey, 2018; Dutta et al., 2021) and the arid Deserts and Xeric Shrublands terrestrial ecoregion of Pakistan (Roberts, 1997). Our model predicts the presence of sup-optimal habitat in central and north-western India that has riverine connections to the highly suitable Terai region. Therefore Fishing Cat populations here could be an extension of the Terai population. However, these could be dead ends as they meet the Thar Desert to the west and the highlands and plateau regions of Central India to the south. Nonetheless they represent threatened freshwater ecosystems worthy of attention. In Pakistan, which is the westernmost limit of global Fishing Cat distribution, it is the presence of the Lower and Middle Indus freshwater ecoregions (see Abell et al., 2008), within the drier terrestrial ecoregion that has made this region highly suitable for Fishing Cat with several priority conservation units. Thus records of Fishing Cat from drier regions suggest that either these are marginal habitat or that climatic and geographic factors have contributed to the formation and persistence of wetlands within the drier regions which facilitated the cat to colonize these regions.

Thirdly, our model predicts Fishing Cat occurrence in the Indian western coast which is dominated by the Western Ghats mountain range and associated streams. There are no confirmed reports of the cat’s occurrence here historically. Recent surveys have been unable to detect the cat in the western coast, following which it was hypothesized that the salinity of the Arabian Sea could have been the limiting factor (Janardhan et al., 2014). We suggest that rather than salinity, the absence of river systems forming extensive floodplains and deltas in this region, is responsible for Fishing Cat’s inability to colonize it. Presence of such riverine systems can overcome limitations imposed by salinity, as is found in the Indus delta region, which meets the Arabian Sea to the south.

Fourthly, our model predicted a gain of 19% habitat between 2010 to 2020 (Table 2). This predicted gain is a result of the significant number of occurrence points in the proximity of urban areas. Such landscapes, which are by-products of rapid urbanization replacing wetland habitat are neither non-habitat, nor fully suitable ones, where the cat has no choice but to persist in remaining small patches. Such landscapes thus exhibit an extremely complicated mosaic of available habitat and built-up spaces, making it difficult for the model to distinguish between pure habitat and mixed/non-habitat, leading to counter-intuitive predictions suggesting habitat gain. For instance, in Colombo, Sri Lanka, GPS-collared adult females were found to use wetland areas more than urban areas whereas a sub-adult male was found to exclusively use hyper-urban areas within the city suggesting that it was pushed into marginal habitat by adults (Ratnayaka et al., 2021). These hyper-urban areas, as observed by us, were characterized by abandoned colonial architectures replete with anthropogenic food sources such as fish present in large courtyard ponds, commensal rodents and birds present in the unkempt gardens. However, all urban spaces, especially upcoming modern smart cities do not/will not have heritage buildings and their unmanicured open spaces, which is why the gain shown by our model cannot be ecologically supported. Moreover, road kills and negative interactions between humans and Fishing Cat are commonly occurring issues in rapidly urbanizing landscapes (Adhya et al., 2011; Thudugala, 2015; Ganguly et al., 2020; Thudugala *in litt*). In Sri Lanka alone, data from a citizen science program (2005 to 2021) shows 101 records of cat death/injury from vehicular collisions. This further suggests that urban spaces might infact be ecological traps in the long term.

### 4.2 Research and conservation efforts needed outside PA network

According to model predictions, more than 90% of the predicted Fishing Cat distribution lies outside protected areas. However, qualitative and/or quantitative assessments of populations and their threats outside protected areas in range countries remain woefully scarce. The few studies that have been conducted on the felid largely report sporadic occurrences and status reports that either lacked rigor, comprehensiveness and/or placement within the larger biogeographic context (Adhya and Dey, 2011; Mukherjee et al., 2012; Palei et al., 2018; Chutipong et al., 2019; Kantimahanti et al. 2019; Kolipaka et al., 2019; Chakraborty et al., 2020). Population estimates on the species are very scarce and restricted to a few protected areas (Nair, 2012; Das et al., 2017) with the exception of Phosri et al., 2021. Across South Asia, it is the freshwater landscapes outside protected areas that seem to be important for maintaining connectivity but are being subjected to severe degradation. Large portions of the Indus, wetlands of the Gangetic Plains, particularly in Bangladesh, the Terai-Dooars region, wetland complexes of the Brahmaputra and coastal wetlands of the Indian eastern coast between Chilika lagoon and the southernmost Indian Fishing Cat population in Godavari-Krishna delta need intensive surveys to not only determine current status but also to examine present connectivity using advanced genetic tools. The Indus population deserves special mention as a region that holds the westernmost population of Fishing Cat globally, is predicted to have highly suitable areas important for conservation, but could well be suffering from isolation effects. Roberts (1977), had reported occasional stragglers from the Middle Indus Basin, and, along the floodplains of the rivers Sutlej, Beas and Ravi in the adjacent Indian Punjab district. A thorough comprehensive current assessment is needed in the region.

The distribution of the species in most of South-east Asia has been extremely hampered in the recent past. We recommend thorough investigations and rapid conservation efforts to stem the loss of low-elevational wetlands like mangroves and marshes in the pinch points of connectivity. The species has been detected in the Ayeyarwady region of Myanmar (Lin and Pratt, 2019). Targeted surveys are required in the adjacent Salween Basin which has moderately suitable habitat. In Thailand, Fishing Cat has been detected in the Chao Phraya river system region (Chutipong et al., 2019) and from peninsular Thailand (Cutter, 2007). More investigations are recommended in the inland and coastal wetlands present in the country’s eastern and western coastal provinces (see S1), especially around the Krabi estuary, even though Chutipong et al., (2019), failed to detect them in this area. In Cambodia, Fishing Cat populations were found to be restricted to the mangrove forests of Peam Krasop Wildlife Sanctuary and Ream National Park but not in Botum Sakor National Park (Thaung et al., 2018). Longer, more intensive and targeted surveys focussed in coastal wetlands of Cambodia as well as along shoreline mangrove patches, for example, in the Koh Kong Krov island are needed. Inland wetlands along the Mekong river in Cambodia should also be targeted for further surveys. A dead specimen retrieved from a roadside market in the Tonle Sap area in 2018, indicates the possibility of an existing population there (Ministry of Agriculture Forestry and Fisheries and Wildlife Alliance *pers. comm.*, 2018) as was previously assumed (Davidson et al., 2006; Rainey and Kong, 2010), prompting surveys currently underway. In Vietnam, where surveys have failed to detect Fishing Cat in recent decades, we recommend targeted surveys in the Plain of Reeds, Red River Delta and U Minh wetlands. Part of U Minh consists of peat swamps which might not be preferred by Fishing Cat but parts with swamps and grasslands could be potentially important survey sites. In Java, there is evidence of the cat’s presence historically (Sody, 1936) and more recently (Duckworth *pers. comm.*, 2012) corroborating model findings. However, surveys have failed to re-detect it since the last decade suggesting that the species is critically endangered in the country (Willianto *pers. comm.*, 2021). No suitable habitat for Fishing Cat was predicted to be present in Sumatra. This is backed by absence of either historical or recent evidence of its presence in the country. How the Fishing Cat came to occur in Java therefore is a mystery.

### 4.3 Loss of critical wetland habitat

Our models predict a contraction of more than 23.74% in suitable Fishing Cat habitat between the two time periods TI and T2 due to urbanization. This is relatable given the continuing trend in rapid wetland loss globally and especially in Asia (Davidson, 2018). Wetlands are predicted to disappear much faster given the projected increase of urban population by 1.4 billion within 2050 (Hettiaracchi et al. 2015). Moreover, ephemeral or intermittently flooded wetlands are not well-represented in global datasets (Davidson, 2014), which are known to be habitat preferred by Fishing Cat, such as reed-building marshes, mangrove swamps and freshwater swamps (Sunquist and Sunquist, 2014; Mukherjee et al., 2016; Hunter et al., 2019). This further augments the threat to Fishing Cat as loss of their primary habitat due to urbanization is undoubtedly rapid but has remained undetected and therefore not foregrounded.

Human populations are known to be concentrated in productive lowland regions of the world and undoubtedly it is here that the impact of sustaining modern industrial human societies are most strongly felt (Darwall and Freyhof, 2016). Coastal wetlands have been more severely affected by urbanization and its indirect effects than inland natural wetlands (Wilkinson et al., 1994; Davidson, 2018). For instance, continuous development pressures have been degenerating the Indo-Malayan mangroves with several stretches in coastal South and South-east Asia annihilated to create megacities (Dasgupta and Shaw, 2013; Kumar and Ambastha, 2018). The loss has exacerbated in the last few decades rendering mangroves in the Indus, Ayeryarwady, Mekong and Red river basins critically degraded with the Lower Indus Basin losing 43% of its mangrove cover from 1990 to 2014 and Ayeyarwady delta losing 64% from 1978 to 2011 with predictions of complete deforestation by 2026 (Dasgupta and Shaw, 2013; Webb et al., 2014; Richard and Freiss, 2016; Qasim et al., 2016). Commercial shrimp culture is a particularly severe threat to the coastal wetlands of South-east Asia destroying 41% of Thailand’s coastal mangroves and 50% of Mekong delta’s mangroves since 1975 (Dasgupta and Shaw, 2013). Wetlands along the Indian east coast formed by Mahanadi, Godavari and Krishna have also been affected by hydrological fragmentation and dams with recent studies estimating a decline in the Godavari-Krishna delta by 42 km^2^ between 1977 and 2008 (Kumar and Patnaik, 2018; Day et al., 2019). Mangroves in this region were found to be under high stress from intensive aquaculture and industrialization (Day et al., 2019; Bagaria et al., 2021). In addition, 42% riparian vegetation has also been lost in the Lower Indus Basin from 1990 to 2014 (Webb et al., 2014). In the Terai Arc Landscape shared between Nepal and India, much of the wet grasslands and swamps were converted during 1970-80 for agricultural expansion and human settlements with only patches of disjunct habitat persisting in protected areas (Chanchani et al., 2014). A further loss of natural grassland area by 24% was estimated from 1980- 2000 across eight protected areas in India and Nepal (Banerjee et al., 2020). In this context, lowlands in GBB which were suggested to be a probable climate refugia for the species by Silva et al. 2020, maybe rendered unsuitable if the present rates and types of alteration continue. According to the Asian Wetland Directory (1989), 68% of Sri Lanka’s 41 internationally significant wetland sites are under moderate to high threat. The Dry Zone’s coastal lagoons, estuaries, and mangrove swamps are highly threatened due to various developmental activities. The Mahaweli Ganga Floodplain System has been recognized as a wetland site ‘under significant threat of destruction’, and is comparable with the Sundarbans and Mekong Delta, demanding immediate action (IUCN Sri Lanka, 2004). In the light of these recent and rapid changes, we further stress the criticality of the status of Fishing Cat worldwide. This is because range reduction is known to imply population declines (IUCN Standards and Petitions Subcommittee, 2017; Ceballos et al., 2017; Santini et al., 2019).

### 4.4 Priority regions and threats

While spatial prioritization for conservation research and action has foregrounded biodiversity in the global context and eased the process of systematic planning and fund acquisition (Myers, 2003), it has also been substantially criticized for being bereft of ‘situated knowledge’ and its failure to engage with locally obtained experience (Wyborn and Evans, 2021). In our study, we attempted to interpret and discuss the findings of our model based on our collated local and regional work experience. Most of the authors are members of Fishing Cat Conservation Alliance, a global cohort of researchers and conservationists, working to facilitate change across the range countries.

Our findings suggest that GBB, consisting of wetlands in the Terai region, Gangetic plains, Gangetic Delta and valleys of the Brahmaputra along with its tributaries and distributaries, have areas with high occurrence probabilities of Fishing Cat (>80%) and might be playing a critical role in maintaining connectivity between populations within South Asia barring the island country of Sri Lanka. The populations in the Terai and Sundarbans were in fact found to be genetically connected in the past (Mukherjee et al., 2015). This implies that populations in GBB are integral to maintaining connectivity between transboundary regions connecting Nepal, Bangladesh and India. Due to its strategic position, GBB is probably indispensable in preventing isolation of populations in the Lower and Middle Indus Basin and those that are located in the Indian East coast populations. Secondly, the Gangetic Delta of GBB is critical in maintaining connectivity between populations in South and South-east Asia, since it is contiguous to the coastal wetlands of Myanmar. Additionally, GBB had the highest number of LCUs, many of which are contiguous, suggesting that this region provides high permeability conducive for species dispersal and requires lesser investments for conservation and restoration of the species’ habitat (Table 3). The Lower Indus Basin had the second highest number of LCUs (Table 3). Even though our connectivity analysis shows that Beas, Ravi and Sutlej floodplains provide limited resistance, this is unlikely given the uninhibited loss of wetlands in the Indian state of Punjab following the Green Revolution and urbanization (Brar and Chandel, 2012). A limitation of our study is the use of only an urbanization based resistance matrix for assessing connectivity, which under-represents the accumulated resistance to dispersal from hostile land-use regimes such as extensively occurring intensive agricultural landscapes and similar aquaculture waterscapes.

The river basins with predicted Fishing Cat occupancy are also among the biggest freshwater fish producing regions with the Ganges, Brahmaputra and Mekong featuring among the top seven basins to contribute to 50% of the global freshwater fish catch, supporting the livelihood of millions locally and providing nutrition globally (WWF, 2021; Ainsworth et al., 2021). These basins are prey-dense wet landscapes for Fishing Cat. Yet these are facing above-average stress levels from the compounding effects of hydropower dams, pollution, overexploitation and climate change resulting in catastrophic declines in wild freshwater fish populations (FAO, 2020; WWF 2021). A third of the world’s freshwater fishes are threatened with extinction and 80 species are already extinct (WWF, 2021). To state that freshwater fishes in the developing Asian economies are being similarly, if not more severely affected, will not be an overstatement. Throughout its global distribution range, 90% of which lies outside the purview of protection, Fishing Cat faces the dual challenge of severe habitat and prey loss. It is here that wetlands are being subjected to diversion to meet development objectives. For instance, in India, these are categorized as ‘wastelands’ (National Remote Sensing Centre, 2010; National Strategic Development Plan, 2014; Hettiaracchi et al. 2015), in contravention of the Ramsar convention and by contradicting national wetlands protection laws (see Wetland Rules, 2020). In Cambodia, on the other hand, regulations can be weakened to develop large swathes of protected areas including mangrove forests to meet economic objectives. The felid was recently downlisted to Vulnerable in 2016 from Endangered in 2010 with the change in the Red List category attributed to improved data quality and not an improved conservation status (Mukherjee et al., 2010; Mukherjee et al., 2016). However, current trends mandate a re-assessment in the light of the species’ critical condition in Java and Vietnam where it remains undetected for the last two decades. In addition, we have most certainly underestimated its range contraction by not accounting for the compounding effects of development-induced intensive resource uses such as agriculture and aquaculture, which as regimes, add to the impacts of urbanization and climate change thereby escalating global threats to its habitat, i.e., freshwater ecosystems.

### 4.5 A flagship mammal to conserve highly threatened freshwater ecosystems

With research and conservation action mostly having a terrestrial and marine focus, the perspective of freshwater ecosystems have been grossly under-represented, impeding fund allocation and contributing indirectly to their high rates of loss (Tickner et al., 2020). Dearth of basic information such as the distribution and status of many freshwater species has resulted in a lack of focussed broadscale conservation plans (Abell et al., 2008). Given the need to highlight the disproportionately threatened and critical status of freshwater biodiversity, Rees et al., 2020, prescribed the adoption of flagship species to promote and motivate public and political engagement. In this context, Fishing Cat was found to have an above average appeal among the general public despite being restricted in its distribution and niche (Macdonald et al., 2017). Devising range-wide strategies to conserve this hypercarnivore and top predator may benefit the ecological community it represents as well as the critical habitat it inhabits, such as ephemeral wetlands (like marshes, mangroves and freshwater swamps) that are situated outside the purview of protection and are highly abused owing to ill-awareness and apathy. We thus propose that Fishing Cat be used as a flagship species to conserve rapidly degrading low-altitude wetlands and their ecological communities in the major river basins of South and South-east Asia. We emphasize that such conservation has to be socio-ecologically sensitive, inclusive of multiple stakeholders and function within a wise-use operating space (see Kumar et al., 2020), knowing well that the dominant terrestrial strategy of ‘fortress conservation’ is inadequate for freshwater biota conservation (Dudgeon et al., 2006).

## Data accessibility

All distributional data needed to evaluate the results and conclusions of the paper are available at www.fishingcat.org.

## Author contributions

Conceptualization - Tiasa Adhya, Partha Dey; Data curation - Priyamvada Bagaria; Formal analysis - Priyamvada Bagaria; Funding acquisition - N/A; Investigation - Tiasa Adhya, Partha Dey, Priyamvada Bagaria; Methodology - Priyamvada Bagaria; Project administration - Tiasa Adhya; Resources - Jim Sanderson; Software - Priyamvada Bagaria; Supervision - Tiasa Adhya, Partha Dey; Validation - Tiasa Adhya, Partha Dey, Anya Ratnayaka, Ashan Thudugala, Vanessa Herranz; Visualization - Tiasa Adhya, Partha Dey, Priyamvada Bagaria; Roles/Writing - original draft - Tiasa Adhya, Priyamvada Bagaria, Partha Dey; Writing - review & editing - Tiasa Adhya, Partha Dey, Vanessa Herranz, Anya Ratnayaka, Ashan Thudugala, Aravind N.A.

## Acknowledgements

We thank all members of the Fishing Cat Conservation Alliance (www.fishingcat.org) who worked over the years in their respective range countries and improved the quality of data available on the Fishing Cat. This was a collaborative venture where the authors voluntarily engaged to develop the manuscript through a series of online meetings during 2019-2021. This research did not receive any specific grant from funding agencies in the public, commercial and/or not-for-profit sectors. We are thankful to Professor A. Townsend Peterson and Geoff Hyde for their valuable suggestions and comments. We are also grateful to Dr Arjun Srivathsa and Dr Ajith Kumar for their specific inputs.

## Appendix Data Sources

| Habitat Variables | Spatial Resolution | Citation |
| --- | --- | --- |
| **Variables used in the Ecological Scenario** |  |  |
| **Worldclim v.2.0 bioclimatic layers (n=19)** | 2.5 minute (~4.6 km) | Fick, S.E. and R.J. Hijmans, 2017. WorldClim 2: new 1km spatial resolution climate surfaces for global land areas. *International Journal of Climatology* 37 (12): 4302-4315. |
| **Monthly mean of Normalised Difference Vegetation Index (NDVI) layers extracted from the 16-Day L3 Global 1 km – MOD13A2v006. Two NDVI layers from the same month were used for calculating the monthly means (n=12).** | 1 km raster layers resampled to 2.5 minute (~4.6 km) | Didan, K. (2015). MOD13A2 MODIS/Terra Vegetation Indices 16-Day L3 Global 1km SIN Grid V006 [Data set]. NASA EOSDIS Land Processes DAAC. Accessed 2020-02-20 from https://doi.org/10.5067/MODIS/MOD13A2.006 |
| **Elevation derived from SRTM GL3v003** | 90 m resampled to 2.5 minute (~4.6 km) | NASA JPL (2013). NASA Shuttle Radar Topography Mission Global 3 arc second [Data set]. NASA EOSDIS Land Processes DAAC. Accessed 2020-07-15 from https://doi.org/10.5067/MEaSUREs/SRTM/SRTMGL3.003 |
| **Terrestrial ecoregions of the World, The Nature Conservancy classification** | Vector data, rasterised at 2.5 minute spatial resolution | Olson, D. M., Dinerstein, E., Wikramanayake, E. D., Burgess, N. D., Powell, G. V. N., Underwood, E. C., D'Amico, J. A., Itoua, I., Strand, H. E., Morrison, J. C., Loucks, C. J., Allnutt, T. F., Ricketts, T. H., Kura, Y., Lamoreux, J. F., Wettengel, W. W., Hedao, P., Kassem, K. R. 2001. Terrestrial ecoregions of the world: a new map of life on Earth. *Bioscience* 51(11):933-938. |
| **Terrestrial ecoregions of the World, Worldwide Fund for Nature classification** | Vector data, rasterised at 2.5 minute spatial resolution | Olson, D. M., Dinerstein, E., Wikramanayake, E. D., Burgess, N. D., Powell, G. V. N., Underwood, E. C., D'Amico, J. A., Itoua, I., Strand, H. E., Morrison, J. C., Loucks, C. J., Allnutt, T. F., Ricketts, T. H., Kura, Y., Lamoreux, J. F., Wettengel, W. W., Hedao, P., Kassem, K. R. 2001. Terrestrial ecoregions of the world: a new map of life on Earth. *Bioscience* 51(11):933-938. |
| **Global Wetlands v3** | 236 m<br>resampled to<br>2.5 minute<br>(~4.6 km) | Gumbricht, T.; Román-Cuesta, R.M.; Verchot, L.V.; Herold, M.; Wittmann, F; Householder, E.; Herold, N.; Murdiyarso, D., 2017, "Tropical and Subtropical Wetlands Distribution version 2", https://doi.org/10.17528/CIFOR/DATA.00058, Center for International Forestry Research (CIFOR), V3, UNF:6:Bc9aFtBpam27aFOCMgW71Q== [fileUNF] |
| **Digital Soil Map of the World** | Vector data,<br>rasterised at<br>2.5 minute<br>spatial<br>resolution | Land and Water Development Division, FAO, Rome |
| **Euclidean Distance from water layer developed using – using the World Water Bodies data released by ESRI Data & Maps** | Raster layer created in ArcGIS with 2.5 minute spatial resolution | ESRI Data & Maps, 2015 |
| **Additional variables for the Urban Impact Scenario** |  |  |
| **Monthly Land Surface Temperature and Emissivity (LST&E) extracted from MOD11C3 Version 6 (n=12)** | 0.05 degree (~5.6 km)<br>resampled to<br>2.5 minute<br>(~4.6 km) | Wan, Z., Hook, S., Hulley, G. (2015). MOD11C3 MODIS/Terra Land Surface Temperature/Emissivity Monthly L3 Global 0.05Deg CMG V006 [Data set]. NASA EOSDIS Land Processes DAAC. Accessed 2020-07-16 from https://doi.org/10.5067/MODIS/MOD11C3.006 |
| **VIIRS Day/Night Band Nighttime Lights Version 1**<br>Tiles with the ranges 00N60E and 75N0E for all the months of 2013 (T1) and 2019 (T2) were downloaded, resampled, rescaled and an annual mean was calculated. | 15 second (~450 m)<br>resampled to<br>2.5 minute<br>(~4.6 km) | C. D. Elvidge, K. Baugh, M. Zhizhin, F. C. Hsu, and T. Ghosh, “VIIRS night-time lights,” International Journal of Remote Sensing, vol. 38, pp. 5860–5879, 2017. |

## Supplementary Information

**S1:** Mainland river basins along with transboundary freshwater units, wetland complexes and jurisdictional boundaries with predicted occurrence of Fishing Cat.

| River basins | Countries | Wetland complexes | Provinces/ States/ Districts |
| --- | --- | --- | --- |
| Indus basin |  |  |  |
| A. Lower Indus Basin | A i) Pakistan | i) Wetland complexes (Haleji lake, Keenjhar lake, Chotiari reserve, Nara canal area, Qualandri creek in the Indus delta, Ketu Bundar mangroves, the Karachi harbour area, Jubho lagoon and Nurri lagoon) | i) Sindh Province |
|  |  | ii) Areas around the Hub dam adjacent to Sindh province and Miani Hor mangroves | ii) Balochistan Province |
| B. Middle Indus Basin | B i) Pakistan | i) Waterlogged and swampy areas formed by seepage from unlined canals | i) Punjab Province |
| Ganges Brahmaputra Basin (GBB) |  |  |  |
| A. Wetlands of the Terai region | A i) India | i) swamps, marshes and seasonally inundated grasslands in National Parks (Jim Corbett, Dudhwa, Katarniaghat, Kishanpur and Valmiki Tiger Reserve) | i) State of Uttar Pradesh |
|  | ii) Nepal | ii) mudflats, reeds beds, freshwater marshes of Koshi Tappu Wildlife Reserve, Jagdishpur Ramsar site, Chitwan National Park, Parsa | ii) Districts of Sunsari, Saptari, Udayapur, Kapilvastu, Kailali |
| B. Wetlands of Gangetic Plains | B i) India | National Park, Ghodaghodi Tal Ramsar site<br>i) shallow wetlands, riverine wetlands and floodplains of Yamuna and Ganga including Okhla Bird Sanctuary, Baghel Tal, Bakhira Lake, Surha Tal, Dahar Lake, Soor Sarovar Bird Sanctuary, Samaspur Lake ; Ramsar site with wetland stretch between Brijghat and Narora; shallow wetlands in Bihar like Kanwar Jheel, Kusheshwarsthan, Bariella, Vikramshila Dolphin Sanctuary, Moti Jheel and various wetland complexes in several districts of West Bengal. | i) States of Delhi, Agra, Uttar Pradesh, Bihar, West Bengal (including districts of Nadia, Murshidabad, Bardhaman, Howrah, Hooghly, East and West Midnapore) |
| C. Wetlands within the valley of Brahmaputra, its tributaries and distributar-ies | C i) India | i) tall grassland wetland habitat, oxbow lakes and riverine wetlands including Deepor Beel Ramsar Site, Majauli island, Dhir-Dilpai-Dakra complex, Tamranga-Dalani complex, Urpod Beel, Chandubi Lake, Orang, Laokhowa, Kaziranga National Park, Jaldapara Wildlife Sanctuary and Gajoldoba wetland | i) State of Assam and West Bengal |
| D. Wetlands of Gangetic Delta | ii) Bangladesh | ii) seasonally flooded marshy wetlands and swamp forests of Haor basin of Sylhet and eastern Mymensingha, Tanguar Haor Ramsar Site, Hakaluki Haor | ii) Districts of Sylhet, Mymensingh, Sunamganj, Moulvibazar |
|  | D i) India | i) seasonally flooded wetlands, depressed lands, | i) State of West Bengal |
|  | ii)<br>Bangladesh | river creeks, mangrove swamps and marshes including the East Kolkata Wetlands Ramsar Site; Sunderbans delta<br>ii) deltaic wetlands including Marijat Haor, Atadanga Haor, Chanda-Bahgia Beel, Sowagram-Gopalpur Beel, Dakatia Beel, Naldanga Beel, Tarail Beel; Sunderbans delta | ii) Districts spanning most of Bangladesh, except for its hilly tracts |
| Mahanadi basin | i) India | i) depressions, floodplain wetlands and ephemeral wetlands including marshes and mangroves connected to Mahanadi, Brahmani and Baitarani rivers; Debi mouth; Bhitarkanika Ramsar site; Chilika lagoon | i) Districts of Kendrapada, Puri, Khordha; Ganjam |
| Godavari-Krishna delta | i) India | i) mangrove swamps, inland and coastal wetlands | i) Districts of Guntur, Krishna, East Godavari, West Godavari, Visakhapatnam, Vizianagaram, Srikakulam |
| A. Ayeyarwady basin<br>B. Salween delta | i) Myanmar<br>i) Myanmar | i) coastal mangroves and freshwater swamp forests<br>i) floodplain wetlands and mangrove forests | i) States of Ayeyarwady, Bago, Rakhine, Yangon, Kachin, Sagaing<br>i) States of Kayin and Mon |
| Chao Phraya river system | i) Thailand | i) riverine wetlands along the Mun, Chi and Songkhram like the Kut Ting Marshland, a Ramsar site; inland and coastal wetlands including Non Thung Thong wetland drained by Tapi river; Bung Khong Long Ramsar Site; Thale sap | i) Provinces of Nongkhai province, Nakohn si Thammarat, Pattani, Narathiwat, Surat Thani, Patthalung, Songkhla, Phang |
|  |  | estuary, Laem Kham marshes; Nong Plak Phraya wetlands, Krabi estuary | Nga, Phuket, Trang, Satun |
| Lower Mekong Basin | i) Cambodia<br>ii) Vietnam | i) flooded forests, swamps, marshes and grasslands including Stung Treng Ramsar site, Tonle Sap Ramsar Site, Boeung Chhmar Ramsar Site, Bassac marshes Ramsar Site<br>ii) Tram Chim National Park or Plain of Reeds Ramsar Site, Lang Sen Wetland Reserve, U Minh wetland complex | i) Provinces of Stung Treng, Battambang, Kampong Thom, Kandal<br>ii) Provinces of Dong Thap, Long An, Kien Giang, Ca Mau |
| Red River Delta | i) Vietnam | i) mangroves and intertidal habitats of Red River Delta Biosphere Reserve | i) Districts of Xuan Thuy and Tien Hai |

**S2:**
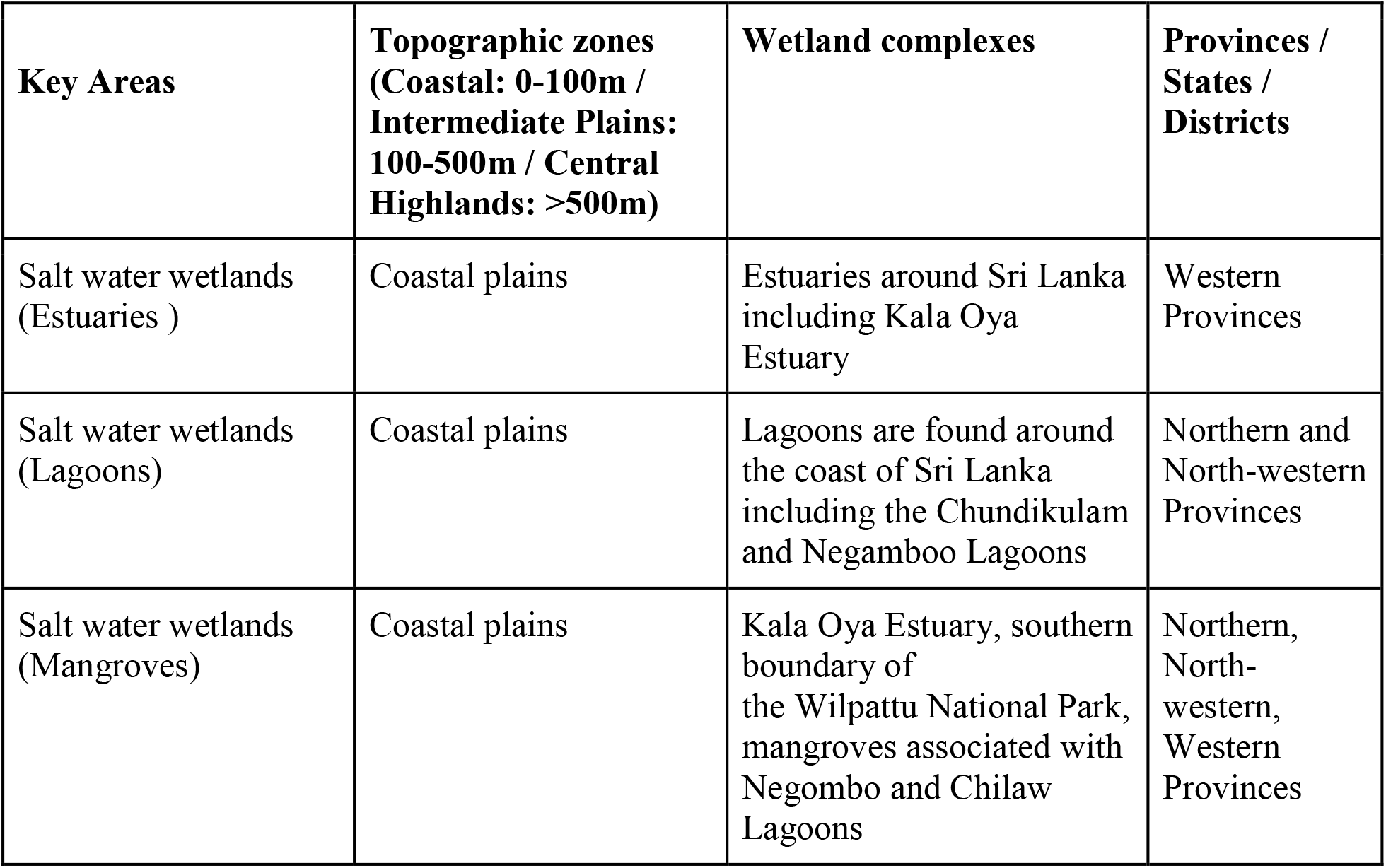
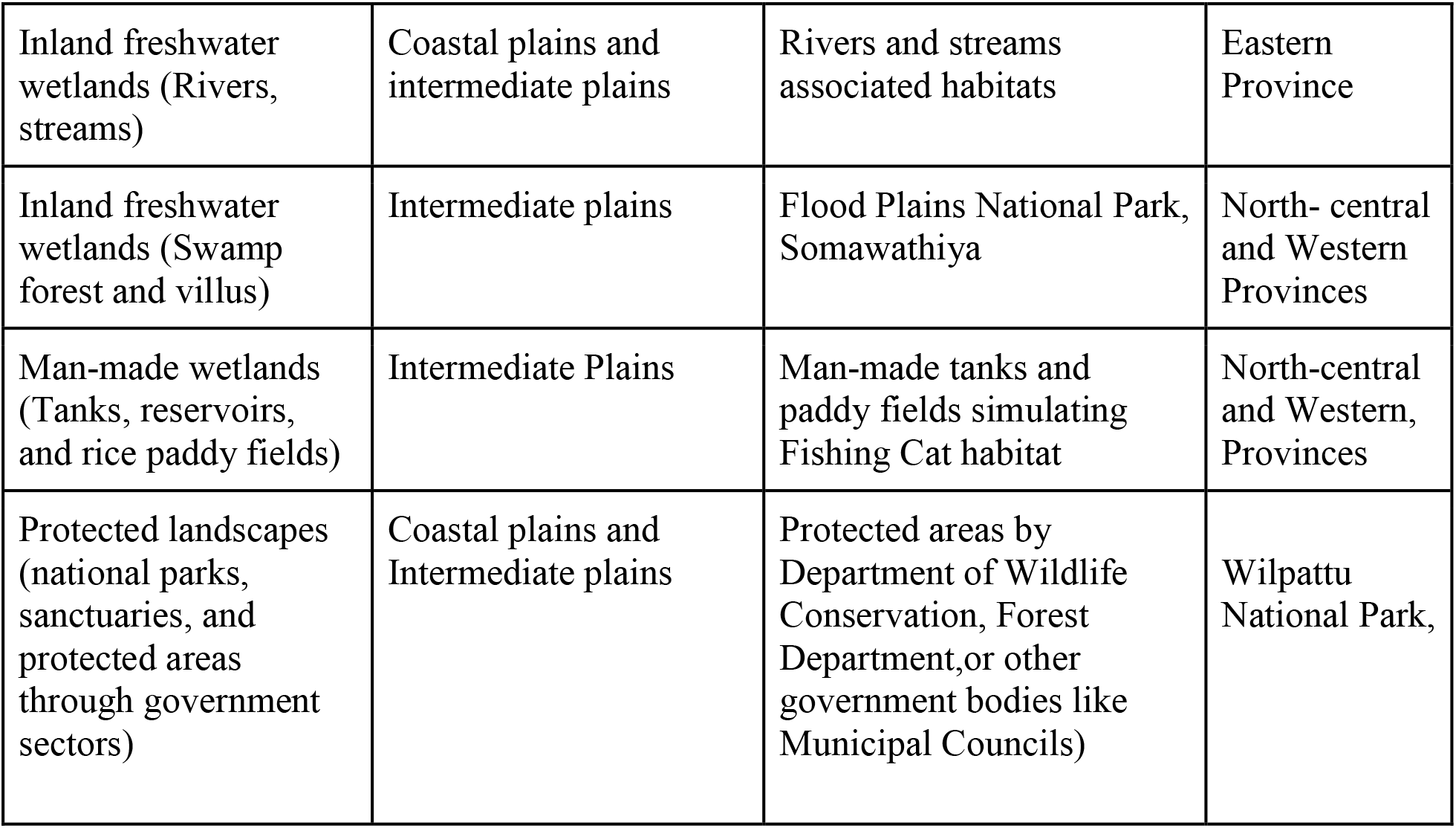
Key areas in Sri Lanka in wetland complexes with jurisdictional boundaries with predicted occurrence of Fishing Cat.

| **Key Areas** | **Topographic zones**<br>**(Coastal: 0-100m /**<br>**Intermediate Plains:**<br>**100-500m / Central**<br>**Highlands: >500m)** | **Wetland complexes** | **Provinces /**<br>**States /**<br>**Districts** |
| --- | --- | --- | --- |
| Salt water wetlands<br>(Estuaries ) | Coastal plains | Estuaries around Sri Lanka including Kala Oya Estuary | Western Provinces |
| Salt water wetlands<br>(Lagoons) | Coastal plains | Lagoons are found around the coast of Sri Lanka including the Chundikulam and Negambo Lagoons | Northern and North-western Provinces |
| Salt water wetlands<br>(Mangroves) | Coastal plains | Kala Oya Estuary, southern boundary of the Wilpattu National Park, mangroves associated with Negombo and Chilaw Lagoons | Northern, North-western, Western Provinces |
| Inland freshwater wetlands (Rivers, streams) | Coastal plains and intermediate plains | Rivers and streams associated habitats | Eastern Province |
| Inland freshwater wetlands (Swamp forest and villus) | Intermediate plains | Flood Plains National Park, Somawathiya | North- central and Western Provinces |
| Man-made wetlands (Tanks, reservoirs, and rice paddy fields) | Intermediate Plains | Man-made tanks and paddy fields simulating Fishing Cat habitat | North-central and Western, Provinces |
| Protected landscapes (national parks, sanctuaries, and protected areas through government sectors) | Coastal plains and Intermediate plains | Protected areas by Department of Wildlife Conservation, Forest Department, or other government bodies like Municipal Councils) | Wilpattu National Park, |

